# Oncogenic RAS commandeers amino acid sensing machinery to aberrantly activate mTORC1 in multiple myeloma

**DOI:** 10.1101/2021.11.28.470260

**Authors:** Yandan Yang, Thomas Oellerich, Ping Chen, Arnold Bolomsky, Michele Ceribelli, Björn Häupl, George W. Wright, James D. Phelan, Da Wei Huang, James W. Lord, Callie K. Van Winkle, Xin Yu, Jan Wisnieski, James Q. Wang, Frances A. Tosto, Erin Beck, Kelli Wilson, Crystal McKnight, Jameson Travers, Carleen Klumpp-Thomas, Grace A. Smith, Stefania Pittaluga, Irina Maric, Dickran Kazandjian, Craig J. Thomas, Ryan M. Young

## Abstract

Oncogenic mutations within the RAS pathway are common in multiple myeloma (MM), an incurable malignancy of plasma cells. However, the mechanisms of pathogenic RAS signaling in this disease remain enigmatic and difficult to inhibit therapeutically. We employed an unbiased proteogenomic approach to dissect RAS signaling in MM by combining genome-wide CRISPR-Cas9 screening with quantitative mass spectrometry focused on RAS biology. We discovered that mutant isoforms of RAS organized a signaling complex with the amino acid transporter, SLC3A2, and MTOR on endolysosomes, which directly activated mTORC1 by co-opting amino acid sensing pathways. MM tumors with high expression of mTORC1-dependent genes were more aggressive and enriched in RAS mutations, and we detected interactions between RAS and MTOR in MM patient tumors harboring mutant RAS isoforms. Inhibition of RAS-dependent mTORC1 activity synergized with MEK and ERK inhibitors to quench pathogenic RAS signaling in MM cells. This study redefines the RAS pathway in MM and provides a mechanistic and rational basis to target this novel mode of RAS signaling.

## Introduction

Multiple myeloma (MM) is the second most common hematological malignancy, accounting for over 32,000 new cancer cases a year within the United States (www.seer.cancer.gov). Substantial progress has been made treating this disease with the introduction of proteasome inhibitors and immunomodulatory drugs (IMiDs). These agents target vulnerabilities tied to the plasmacytic origins of MM and have significantly extended patient survival (*1*, *2*). However, MM remains incurable and most patients will relapse and become refractory to existing treatments. Mutations targeting the RAS pathway are common in MM and associated with resistance to these therapies (*3*). KRAS and NRAS are each mutated in about 20% of newly diagnosed MM cases (*4*, *5*). MM is unusual in this regard, as other RAS-dependent tumor types typically rely on a single isoform of RAS (*6*). RAS can signal through a number of effector pathways, perhaps most characteristically by activation of the classical MAP kinase (MAPK) pathway through RAF, MEK and ERK. Despite the high frequency of RAS mutations, the majority of MM tumors harboring RAS mutations have no detectable MEK activity by immunohistochemistry staining (*7*) or analysis of MAPK-dependent transcription (*8*), and MEK inhibitors have only had modest success treating MM patients in the clinic (*9*, *10*). These findings suggest that RAS-dependent activation of the classical MAPK pathway is not the sole mode of RAS signaling in malignant plasma cells and point to an unidentified role for oncogenic RAS signaling in this disease.

To uncover mechanisms of pathogenic RAS signaling in MM, we implemented an unbiased proteogenomic pipeline that combined CRISPR-Cas9 screens to identify genes selectively essential in MM lines dependent on KRAS or NRAS expression, as well as quantitative mass spectrometry (MS) to determine protein interaction partners for mutant RAS isoforms in MM cells. This approach revealed the “essential interactome” of mutant RAS and highlighted the connection between RAS and SLC3A2. SLC3A2 (CD98, 4F2hc) is a component of several heterodimeric amino acid transporters for large neutral amino acids, including SLC3A2-SLC7A5 that serves to transport leucine and glutamine (*11*). Proteomic analysis of RAS and SLC3A2 interaction partners and dependent signaling networks identified that mTORC1 was activated downstream of both RAS and SLC3A2. We determined that RAS coordinated the co-localization of SLC3A2 and MTOR on LAMP1+ endolysosomes, where RAS, SLC3A2 and MTOR cooperatively activated mTORC1. RAS accomplished this by subverting nutrient sensing pathways that normally regulate homeostasis through mTORC1. Inhibition of RAS-dependent mTORC1 activity enhanced reliance reliance on MEK and ERK signaling in MM cells, and combinations of mTORC1 and MEK inhibitors resulted in a synthetic lethal phenotype that was profoundly toxic to RAS-dependent malignant cells. Thus, our work details a new concept in pathogenic RAS signaling and outlines potential therapeutic opportunities to exploit this novel signaling mechanism.

## Results

### Proteogenomic screens in MM

We conducted CRISPR-Cas9 screens to identify genes essential to malignant growth and survival in 17 MM cell lines (Fig. 1A). To maximize the sensitivity and utility of these CRISPR screens, Cas9-engineered MM lines were first selected for exceptional exonuclease activity as determined by reduction in CD54 levels following expression of a CD54-targeted single guide RNA (sgRNA) (Fig. 1A). Cas9 clones with high knockout efficiency were subsequently screened with the third-generation genome-wide Brunello sgRNA library (*12*) to identify essential genes after 21 days of growth. For each gene we determined the CRISPR screen score (CSS), a metric of how deletion of a gene affects cell growth and survival, akin to a Z-score (*13*) (Table S1). Deletion of genes known to be essential to MM biology, including XPO1 (*14*), IRF4 (*15*), MYC (*16*) and MCL1 (*17*) were toxic to all MM lines and had negative CSS values (Fig. S1A). In contrast, FAM46C and ID2 acted as tumor suppressors in many MM lines as indicated by their positive CSS values (Fig. S1A), which is consistent with previous results (*18*, *19*). Comparison of our CRISPR dataset to the 20 MM cell lines within DepMap (*20*) found general agreement (Fig. S1B), although our approach identified additional MM-specific essential genes, likely due to the superior performance of the Brunello library (*21*).

**Figure 1.**
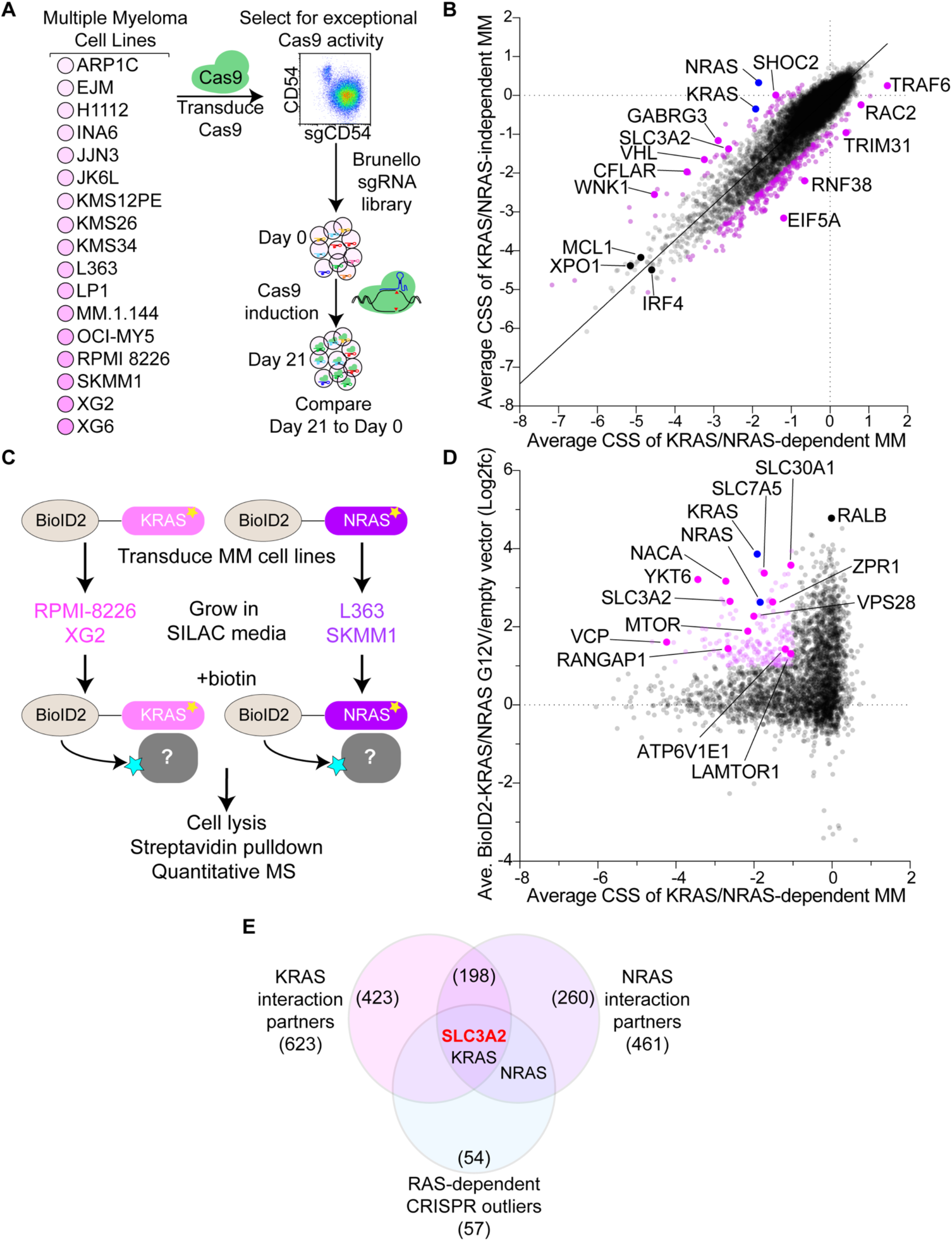
Proteogenomic screens reveal SLC3A2 is a novel RAS effector in multiple myeloma. (A) Workflow for CRISPR screens in MM cell lines. (B) Scatter plot of the average CSS for RAS-dependent MM lines (x-axis) vs. RAS-independent MM lines (y-axis). Outliers were determined by an extra sum-of-squares F test (p=0.02) and are labeled in purple. (C) Workflow of BioID2-based KRAS and NRAS interactome SILAC MS studies. (D) The essential interactome of G12V RAS isoforms in RAS-dependent MM. The average CSS for KRAS/NRAS-dependent MM cell lines (x-axis) plotted by average combined enrichment of BioID2-KRAS^G12V^/ BioID2-KRAS^G12V^ relative to empty vector. The essential interactome (≤-1.0 CSS and ≥1.0 log2fc BioID2-RAS) is labeled in pink. (E) Venn diagrams of protein interaction partners enriched by ≥2.0 log2fc in BioID2-KRAS^G12V^ and BioID2-NRAS^G12V^ with RAS-dependent outlier genes from Fig. 1B.

We next parsed the CRISPR screen data to identify genes preferentially essential in RAS-dependent MM. Our screens identified eight MM cell lines reliant on KRAS or NRAS expression for their growth and survival (Fig. S1C). All RAS-dependent MM lines harbored oncogenic RAS mutations, with the exception of KMS26 which expressed wild type KRAS. In contrast, EJM, JJN3, JK6L and XG6 expressed mutant isoforms of RAS but were not sensitive to KRAS or NRAS knockout. We compared CRISPR screen results from RAS-dependent MM lines (grouped KRAS-dependent and NRAS-dependent, x-axis) to MM lines insensitive to KRAS or NRAS deletion (y-axis) in Figure 1B. Outlier genes selectively more toxic in RAS-dependent or RAS-independent MM lines were identified by an extra sum-of-squares F test (p<0.05) (Fig. 1B; Table S2). In addition to NRAS and KRAS themselves, the RAS-dependent outliers included SHOC2, which was previously shown to activate MAP kinase (MAPK) signaling downstream of oncogenic RAS (*22*). However, most other RAS-dependent outliers have no reported link to RAS signaling, and pathway analysis of these genes yielded no significant enrichments or clues to their function.

To unlock additional insight from these CRISPR screens, we employed an orthogonal proteomic approach to identify proteins that interact with RAS isoforms in MM cells. BioID2 is a promiscuous biotin ligase that can biotinylate proteins within a 10-30 nm distance (*23*), and we ectopically expressed BioID2 fused to KRAS^G12V^ in RPMI 8226 and XG2, or NRAS^G12V^ in SKMM1 and L363 MM cells (Fig. 1C). Biotinylated proteins were purified from these cells by streptavidin pulldown and enumerated using quantitative stable isotope labeling by amino acids in cell culture mass spectrometry (SILAC-MS) (Fig. 1C). These experiments identified numerous proteins enriched relative to empty vector vLYT2-BioID2 expressing BioID2 alone (Fig. S2A-B), including many known RAS effectors (Fig. S2C; Table S3). To focus on the RAS interactors most essential to growth and survival in MM, we compared the enrichment of proteins within BioID2-RAS interactomes to the CRISPR screen data, both as an average of all RAS-dependent MM cells (Fig. 1D). Notably, these essential interactomes did not include classical RAS effectors – including BRAF and RALA – because these genes were not determined to be essential by CRISPR screening, although it remains possible this is due to redundancy among paralogous genes. Regardless, this essential interactome highlighted associations between RAS and MTOR, several solute-carrier (SLC) genes, and many cellular trafficking proteins. Shared KRAS and NRAS interaction partners were significantly enriched in Gene Ontology (GO) pathway gene sets (*24*) associated with membrane and vesicular trafficking (Fig. S2D). We next compared KRAS and NRAS protein interaction partners (≥2 log2fc) with the set of genes found to be significantly more essential within RAS-dependent tumors (Fig. 1B). Remarkably, only SLC3A2 interacted with both KRAS and NRAS, and was more selectively essential in RAS-dependent MM lines (Fig. 1E).

### SLC3A2 regulates mTORC1 signaling in RAS-dependent MM cells

SLC3A2 (CD98, 4F2hc) is integral to amino acid transport into cells (*11*), and high levels of expression are correlated to aggressive MM (*25*). We confirmed that SLC3A2 associated with RAS isoforms in MM cells by co-immunoprecipitation with ectopically expressed mutant isoforms of KRAS or NRAS in various MM cell lines (Fig. 2A). To explore SLC3A2 function in MM cells, we expressed a BioID2-SLC3A2 construct in RPMI 8226 to determine SLC3A2 protein interaction partners by MS. We resolved the SLC3A2 essential interactome by plotting the BioID2-SLC3A2 protein enrichment (y-axis) against CSS values for RPMI 8226 (x-axis) (Fig. 2B). As expected, we observed a strong interaction between KRAS and SLC3A2 in RPMI 8226 cells. In addition, we detected robust interactions with other essential and non-essential SLC-family genes, including SLC7A5, SLC38A1, SLC30A5 and SLC4A7. SLC3A2 is known to heterodimerize with SLC7A5 (*26*), but many of these SLC genes have not been previously described to interact with SLC3A2 and it is unclear if all these associations represented novel heterodimers with SLC3A2 or perhaps reflected the proximity of these proteins in the cell membrane. However, our BioID2 data suggested that SLC3A2 primarily pairs with SLC7A5 to form a leucine and glutamine transporter (*26*).

**Figure 2.**
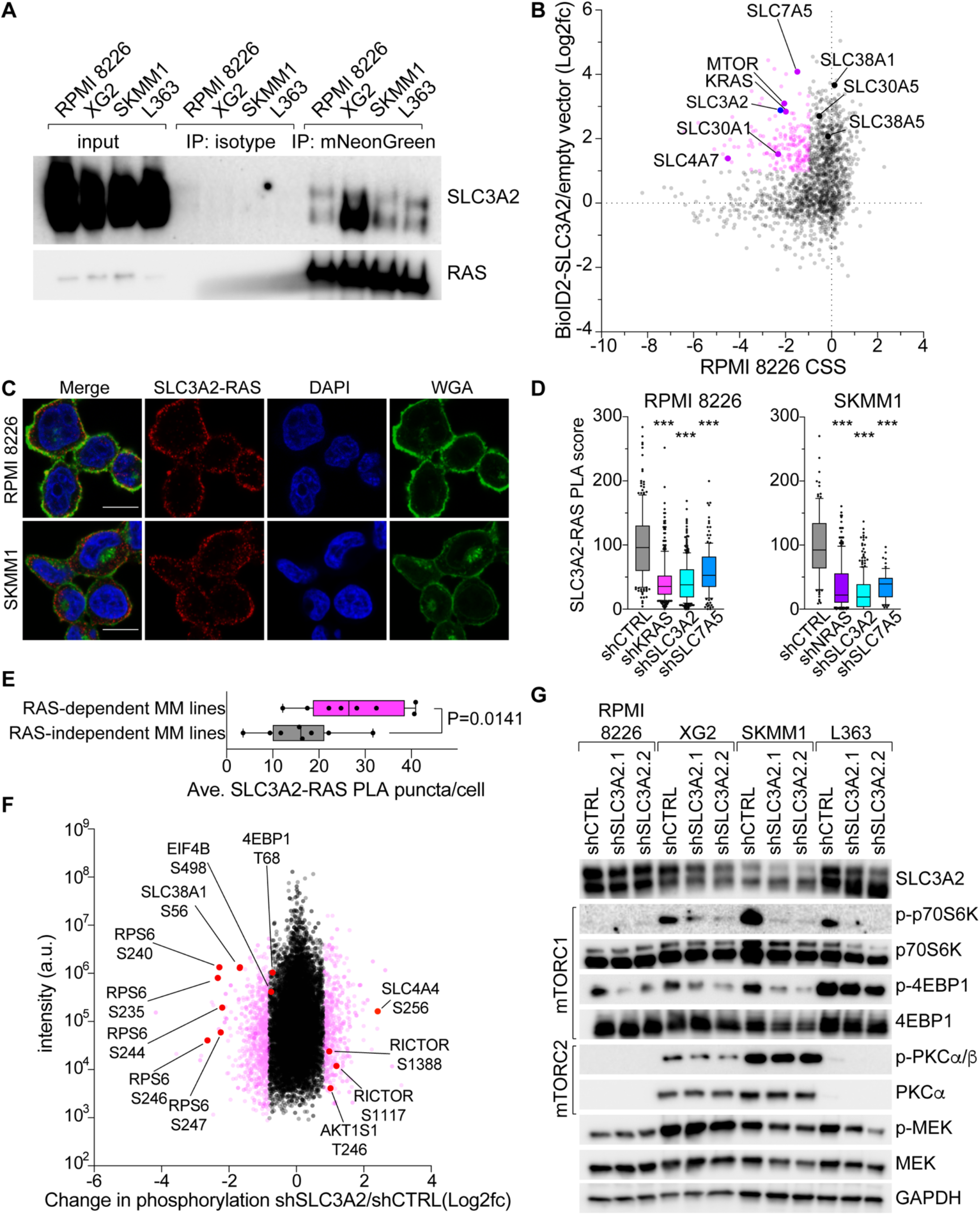
SLC3A2 interacts with RAS and controls mTORC1 activity in MM. (A) Co-immunoprecipitation of SLC3A2 with mutant isoforms of mNeonGreen-tagged KRAS and NRAS. mNeonGreen-tagged KRAS^G12D^ was used in RPMI 8226 and XG2, NRAS^G12D^ in SKMM1 and NRAS^Q61L^ in L363. Representative blots from 4 independent experiments. (B) The essential interactome of SLC3A2 in RPMI 8226. The CSS (x-axis) plotted by the BioID2-SLC3A2/empty vector enrichment (y-axis). The essential interactome is labeled in pink with callouts in red. (C) PLA of SLC3A2 and RAS in RPMI 8226 and SKMM1 cells. PLA is shown in red, DAPI in blue and WGA in green. Scale bar is 10μm. Representative images from 3 or more experiments. (D) PLA score of cells transduced with control shRNA or shRNAs specific for KRAS, NRAS, SLC3A2 and SLC7A5. *** denotes p-value <0.0001 by one-way ANOVA, values from 3 or more experiments. (E) The average SLC3A2-RAS PLA puncta/cell for 16 MM cell lines. Each dot represents an individual MM line. P-value from one-tailed Mann-Whitney test. Representative data from two independent experiments. (F) Global changes in phosphorylation measured by quantitative MS following knockdown of SLC3A2 in SKMM1. Change in phosphorylation (log2fc) in cells expressing shSLC3A2 vs. shCTRL (x-axis) plotted by the measured intensity (y-axis). (G) Western blot analysis of mTORC1 and mTORC2 effectors in MM lines transduced with shCTRL, or RAS-specific shRNAs. Representative blots from 4 experiments.

We next used the proximity ligation assay (PLA), which can quantitatively visualize proteinprotein interactions within tens of nanometers as discrete puncta *in situ* (*27*), to study interactions between endogenous SLC3A2 and RAS in the KRAS-dependent RPMI 8226 and NRAS-dependent SKMM1 MM cells. We observed numerous bright PLA puncta confirming the proximity of SLC3A2 and RAS in both MM lines (Fig. 2C; red). We found PLA signal near the plasma membrane, stained by wheat germ agglutinin (WGA; green), as well as within the cytosol. Immunofluorescence staining of SLC3A2 and RAS showed that both proteins are predominantly localized to the plasma membrane but also share a diffuse staining throughout the cytoplasm that was highlighted by PLA (Fig. S3A). Knockdown of RAS isoforms or SLC3A2 abrogated SLC3A2-RAS PLA signal, demonstrating the specificity of detecting this interaction by PLA (Fig. 2D). Knockdown of SLC7A5 also abolished PLA signal between SLC3A2 and RAS (Fig. 2D), providing more evidence that SLC3A2 likely interacts with RAS as part of a heterodimer with SLC7A5, as supported by our proteomic data (Fig. 2B). Finally, analysis of SLC3A2-RAS PLA in the cohort of MM lines used for CRISPR screening found that RAS-dependent MM cell lines had significantly more PLA puncta per cell than RAS-independent MM cell lines (Fig. 2E).

We noted that SLC3A2 strongly interacted with MTOR in BioID2 proximity labeling experiments (Fig. 2B). MTOR regulates cellular growth, metabolism and proliferation as a member of two multicomponent signaling complexes, mTORC1 and mTORC2. mTORC1 signaling is gated by the availability of nutrients, such as amino acids, and SLC3A2 has been previously implicated in its regulation in its role as an amino acid transporter (*26*). We probed the role of SLC3A2 in MM signaling by enumerating changes in global phosphorylation by quantitative MS following SLC3A2 knockdown in SKMM1 cells (Fig. 2F; Table S5). SLC3A2 knockdown substantially decreased phosphorylation of RPS6 at multiple serine residues. RPS6 is a target of p70S6K, a known effector downstream of mTORC1 (*28*). Western blot analysis following SLC3A2 knockdown in RAS-dependent MM lines found significant reductions in mTORC1 targets (p-p70S6K (T389) and p-4EBP1 (S65)) but minimal changes in the mTORC2 target PKCa (p-T638/641) (Fig. 2G). These data confirmed that SLC3A2 regulated mTORC1 signaling in MM and demonstrated that RAS was required for this activity. Of note, phosphorylation of MEK, a target of RAS signaling, was only modestly reduced by SLC3A2 knockdown, suggesting that although SLC3A2 and RAS are interaction partners, SLC3A2 does not substantially control MEK signaling downstream of RAS (Fig. 2G).

### RAS controls association of SLC3A2 with MTOR on LAMP1+ endolysosomes

To understand how RAS may regulate SLC3A2 in MM cells, we evaluated changes in SLC3A2 protein interaction partners following RAS knockdown. We expressed BioID2-SLC3A2 in four RAS-dependent MM lines and enumerated changes in protein interactions by SILAC MS two days after induction of either a control shRNA or shRNAs targeting KRAS or NRAS, corresponding to the mutant isoform of RAS expressed in each MM cell line. Data for the KRAS-dependent MM line XG2 is shown in Figure 3A, and protein interactions that either decreased or increased by 0.5 log2fc are depicted in purple or blue, respectively, with outliers labeled if they were found in two or more MM lines (Table S4). We found that SLC3A2 association with MTOR consistently decreased following RAS knockdown. We next used PLA to visualize interactions between endogenous MTOR and SLC3A2, which generated bright puncta throughout the cytosol (Fig. 3B, red). The MTOR-SLC3A2 PLA was specific since knockdown of its constituent parts, MTOR and SLC3A2, nearly eliminated PLA signal (Fig. 3C). In addition, knockdown of KRAS or NRAS substantially reduced interactions between MTOR and SLC3A2, confirming the BioID2 results in Figure 3A and demonstrating that RAS governs associations between SLC3A2 with MTOR.

**Figure 3.**
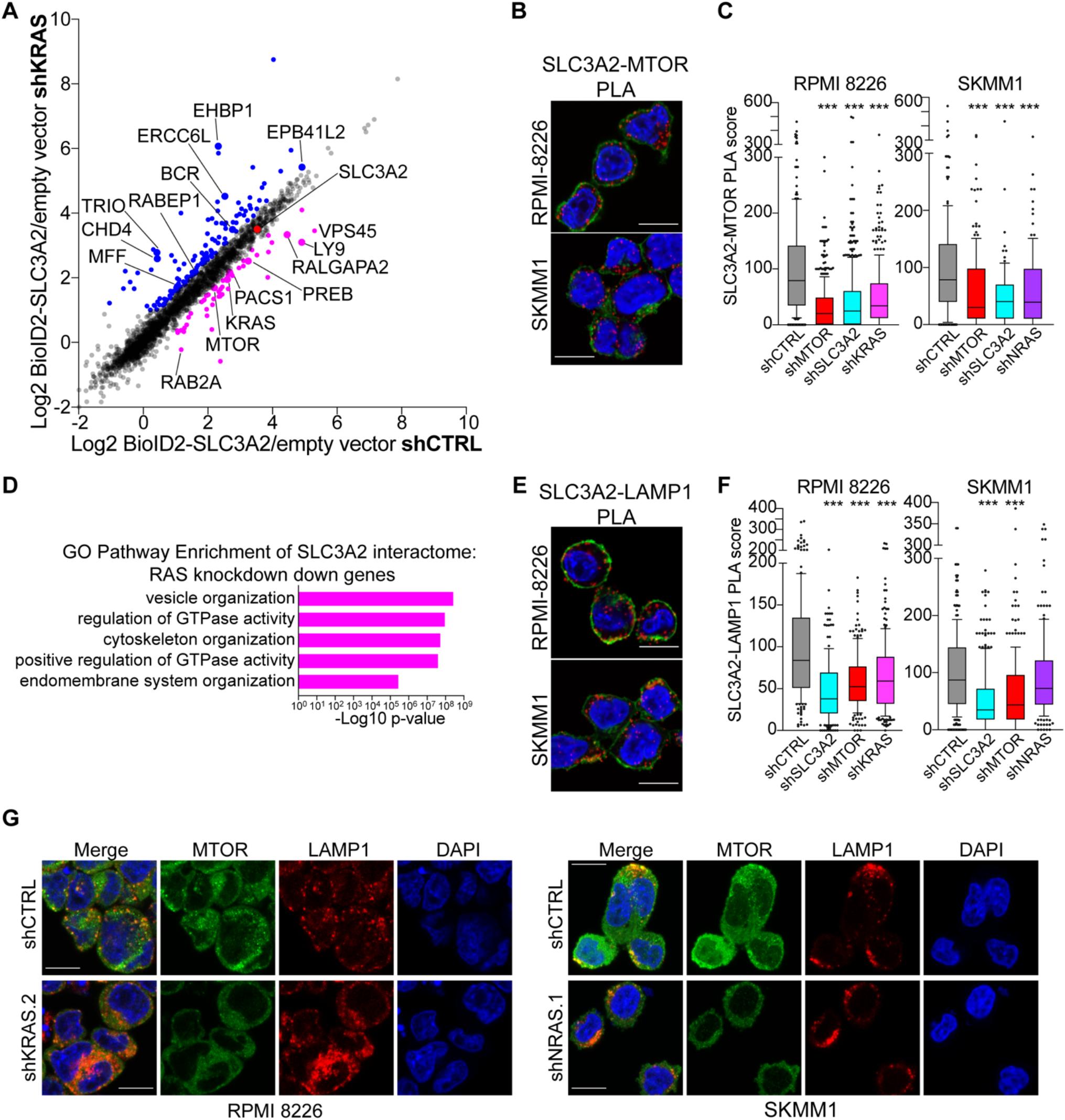
RAS regulates intracellular trafficking of SLC3A2 to endolysosomes and MTOR. (A) BioID2-SLC3A2 interactomes in XG2 cells transduced with control shRNA (x-axis) or shKRAS (y-axis). Proteins that decreased by 0.5 log2 fold-change or more are labeled in purple, while proteins that increased by 0.5 log2 fold-change or more are labeled in blue. Callouts are for proteins that showed up in at least 2 out of 4 MS experiments. (B) PLA of SLC3A2 with MTOR in RPMI 8226 and SKM11 cells. Scale bar is 10μm. PLA is shown in red, DAPI in blue and WGA in green. (C) PLA score of cells transduced with control shRNA or shRNAs specific for KRAS, NRAS, SLC3A2, SLC7A5 and MTOR. *** denotes p-value <0.0001 by one-way ANOVA. All images representative of at least 3 experiments. (D) Gene Ontology pathway enrichment or SLC3A2 interactors that decreased following RAS knockdown. Bonferroni corrected p-value plotted on the x-axis. (E) PLA of SLC3A2 with LAMP1 in RPMI 8226 and SKM11 cells. Scale bar is 10μm. PLA is shown in red, DAPI in blue and WGA in green. (F) PLA score of cells transduced with control shRNA or shRNAs specific for KRAS, NRAS, SLC3A2, SLC7A5 and MTOR. *** denotes p-value <0.0001 by one-way ANOVA. (G) Immunofluorecense images of MTOR (green), LAMP1 (red) and DAPI (blue) in RPMI 8226 and SKMM1 cells expressing shCTRL, shKRAS.2 or shNRAS.1. Scale bar is 10μm. Representative images from 3 independent experiments.

To gain further insight into the mechanisms by which RAS regulates SLC3A2 and MTOR associations, we performed GO pathway analysis on BioID2-SLC3A2 interactors that were reduced by at least 0.5 log2fc following RAS knockdown in any of the four MM lines tested. Proteins with reduced SLC3A2 association following RAS knockdown were enriched in pathways associated with vesicle and endomembrane organization (Fig. 3D), suggesting that RAS may control localization of SLC3A2 to endomembranes. SLC3A2-SLC7A5 has been previously characterized on lysosomal membranes, where it promoted entry of leucine into the lysosomal lumen to stimulate V-ATPase and mTORC1 activity (*29*). To test if RAS regulated trafficking of SLC3A2 to endolysosomes, we developed a PLA pair between SLC3A2 and a marker of endolysosomes, LAMP1. SLC3A2-LAMP1 PLA puncta were observed throughout the cytosol in RPMI 8226 and SKMM1 MM cells (Fig. 3E, red). This interaction was specific, since knockdown of SLC3A2 and MTOR significantly disrupted SLC3A2-LAMP1 PLA signal (Fig. 3F). However, while RAS knockdown reduced the number of SLC3A2-LAMP1 PLA puncta in both MM lines tested, it was only significantly reduced in RPMI 8226 (Fig. 3F). Accordingly, RAS knockdown did not meaningfully change surface expression of SLC3A2 (Fig. S3B), suggesting that RAS does not regulate SLC3A2 trafficking. Alternatively, RAS may control localization of MTOR to endolysosomal membranes in order to bring SLC3A2 and MTOR together on endolysosomes. Immunofluorescence of MTOR and LAMP1 in RPMI 8226 and SKMM1 revealed that MTOR formed foci throughout the cytosol which overlapped with LAMP1 staining (Fig. 3G), consistent with MTOR being engaged in chronic mTORC1 signaling. Knockdown of KRAS or NRAS substantially diminished these MTOR foci, although LAMP1 staining was not reduced (Fig. 3G). Taken together, these data suggest that RAS controls association of SLC3A2 with MTOR by regulating localization of MTOR to LAMP1+ endolysosomes.

### mTORC1 activity is a predominant feature of RAS signaling in MM

Our data support a model where RAS controls mTORC1 activity by coordinating SLC3A2 and MTOR association. This model would predict that RAS regulates mTORC1 signaling in MM. To obtain an unbiased view of RAS signaling in a malignant plasma cell, we employed quantitative MS to enumerate changes in global phosphorylation following knockdown of NRAS in SKMM1 cells. Pathway enrichment of proteins whose phosphorylation changed significantly (+/- 0.8 log2fc) identified that the mTORC1 pathway and, to a lesser extent, the MAPK pathway, are the prominent RAS effector pathways in SKMM1 cells (Fig. 4A; Table S6). NRAS knockdown markedly decreased phosphorylation on targets of mTORC1 signaling (4EBP1, EIF4G1, ULK1) and MAPK signaling (RAF1, MAPK1, MAPK3). In contrast, NRAS knockdown resulted in increased phosphorylation of mTORC2 components and its downstream signaling effectors (MAPKAP1, AKT1, PRKCA) (Fig. 4B), perhaps due to compensatory signaling feedback between mTORC1 and mTORC2. We confirmed these proteomic findings by western blot analysis in additional RAS-dependent MM cell lines. KRAS or NRAS knockdown decreased phosphorylation of mTORC1 targets, p70S6K (T389) and 4EBP1 (S65), in all RAS-dependent MM lines tested (Fig. 4C). We observed little or no change in mTORC1 signaling upon RAS knockdown in MM cells not dependent RAS expression (Fig. S4A). Moreover, disruption of RAS expression also reduced phosphorylation of MEK (S217/221) in these MM lines (Fig. 4C), consistent with the MS phosphoproteomic analysis.

**Figure 4.**
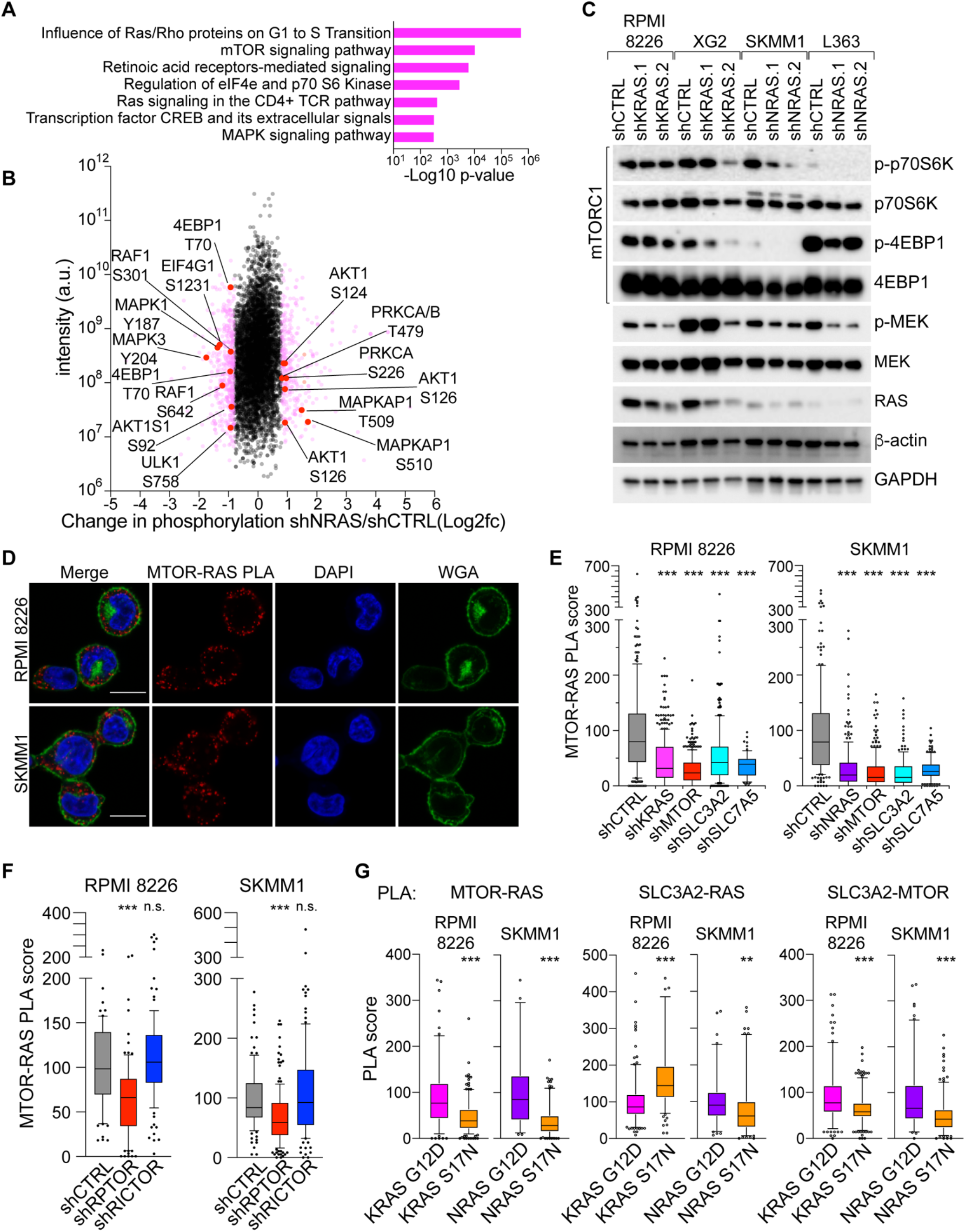
RAS controls mTORC1 activity in MM. (A) Pathway analysis of proteins with +/- 0.8 log2 fold changes in phosphorylation in SKMM1 cells transduced with shNRAS compared to control shRNA as determined by quantitative MS. (B) Scatter plot of changes in phosphorylation in shNRAS/shCTRL (x-axis) vs. intensity (y-axis). Proteins in the mTOR and MAPK signaling pathways are highlighted. (C) Western blot analysis of mTOCR1 and MEK signaling following KRAS or NRAS knockdown in the indicated MM lines. Representative blots from 3 independent experiments. (D) PLA of MTOR-RAS in RPMI 8226 and SKM11 cells. Scale bar is 10μm. PLA is shown in red, DAPI in blue and WGA in green. Representative images from 5 experiments. (E) PLA score of cells transduced with control shRNA or shRNAs specific for KRAS, NRAS, SLC3A2 and MTOR. *** denotes p-value <0.0001 by one-way ANOVA. Data from 5 experiments. (F) MTOR-RAS PLA in RPMI 8226 and SKMM1 cells expressing shCTRL, shRPTOR or shRICTOR shRNAs. *** denotes p-value <0.0001 by one-way ANOVA. Data from 3 experiments. (G) Quantitation of indicated PLAs in RPMI 8226 and SKMM1 cells expressing constitutively active (G12D) or dominant negative (S17N) versions of KRAS or NRAS. *** denotes p-value <0.0001, ** = 0.0028 by Mann Whitney t test. Data from 3 or more experiments.

Our BioID2 studies identified that RAS also interacted with MTOR (Fig. 1D), suggesting that RAS and SLC3A2 may regulate mTORC1 signaling as part of a complex with MTOR. We confirmed that endogenous MTOR and RAS associated in RPMI 8226 and SKMM1 MM lines by PLA (Fig. 4D, red). MTOR-RAS PLA puncta were cytosolic and generally not coincident with the plasma membrane (Fig. 4D, green), consistent with localization to mTORC1 complexes. Knockdown of MTOR and RAS isoforms quenched PLA signals and confirmed the specificity of this PLA pair (Fig. 4E). Additionally, we found that MTOR-RAS PLA signal was significantly enriched in RAS-dependent versus RAS-independent MM cell lines (Fig. S4B). Remarkably, SLC3A2 and SLC7A5 knockdown abolished MTOR-RAS PLA signal (Fig. 4E), suggesting that RAS can only interact with MTOR in the presence of SLC3A2-SLC7A5. MTOR-RAS PLA signal was also significantly correlated to both SLC3A2-RAS PLA signal (Fig. S4C) and the SLC3A2 CSS (Fig. S4D), further linking SLC3A2 with MTOR-RAS associations.

A previous study described that oncogenic RAS isoforms directly bound and activated mTORC2 in melanoma and other solid tumor cell lines (*30*). In contrast, our data support an exclusive role for RAS-dependent activation of mTORC1. To directly test the role of mTORC1 and mTORC2, we assessed whether associations between RAS with MTOR were altered by knockdown of RPTOR or RICTOR, components specific to either mTORC1 or mTORC2, respectively. We found that knockdown of RPTOR substantially decreased the number of MTOR-RAS PLA puncta in RPMI 8226 and SKMM1 cells (Fig. 4F). In contrast, RICTOR knockdown had no effect on MTOR-RAS PLA (Fig. 4F). Notably, we found that RPTOR knockdown also reduced PLA between SLC3A2 and RAS in both RPMI 8226 and SKMM1 (Fig. S5A), demonstrating that mTORC1 expression is required for this association. We also observed robust interactions between RPTOR and RAS in MM cells, but failed to find associations between RICTOR and RAS (Fig. S5B). These data confirm that RAS associates with mTORC1 within MM cells and establishes that mTORC1 signaling is a central feature of RAS and SLC3A2-dependent signaling in MM.

We next sought to understand the role of RAS activity in regulating molecular associations between SLC3A2, MTOR and RAS. RPMI 8226 and SKMM1 cells were transduced with either constitutively active (G12D) or dominant negative (S17N) versions of KRAS or NRAS, respectively. We then evaluated these cells by PLA to measure associations between RAS, MTOR and SLC3A2. We found that expression of dominant negative RAS substantially reduced MTOR-RAS and SLC3A2-MTOR PLA signal in both MM lines (Fig. 4G), suggesting that these protein associations are dependent on RAS activity. In contrast, we observed variable changes on SLC3A2-RAS PLA in MM cells expressing constitutively active or dominant negative RAS isoforms (Fig. 4G), and we cannot conclude if RAS activity is necessary for the association of RAS with SLC3A2. However, these data are consistent with a requirement of RAS activity for localization of RAS, SLC3A2 and MTOR to mTORC1 complexes on endolysosomes.

### Oncogenic RAS co-opts amino acid sensing to activate mTORC1

Our proteogenomic screens illustrated a profound connection between RAS and MTOR signaling in MM. A schematic of MTOR signaling is shown Figure 5A with individual components shaded by their average CSS for RAS-dependent (pink) and RAS-independent (purple) cell lines. Proteins enriched in oncogenic KRAS and NRAS BioID2 experiments by an average of ≥2 log2fc over empty vector are marked by a cyan circle with an ‘R’. This map shows interactions between RAS, SLC3A2, SLC7A5 and MTOR that we have characterized above. Interestingly, genes that comprise mTORC1 were highly essential in all MM cells yet components of mTORC2 were only necessary in RAS-independent MM cell lines, suggesting that RAS-independent MM lines rely on upstream growth factor or chemokine receptor signaling to stimulate phosphoinositide 3-kinase (PI3-K). Nonetheless, we found that oncogenic RAS strongly interacted with components of mTORC2 in BioID2 experiments, although we were unable to confirm associations between RAS and RICTOR by PLA (Fig. S5B). These data raise the possibility that RAS may act to suppress mTORC2 signaling in MM, in accord with changes to global protein phosphorylation following RAS knockdown (Fig. 4B). In addition, Figure 5A shows that RAS associated with numerous components of MTOR signaling in BioID2 experiments, including members of the v-ATPase complex, LAMTOR3, RRAGC and ARF1. We used PLA to visualize interactions of RAS with RAGC, a component of the Ragulator, and ARF1, which can act as a glutamine sensor to activate mTORC1 (*31*) (Fig. 5B). Moreover, we could detect RAS in close association with p-S65-4EBP1 by PLA, suggesting that RAS is at the site of active mTORC1 signaling (Fig. 5B).

**Figure 5.**
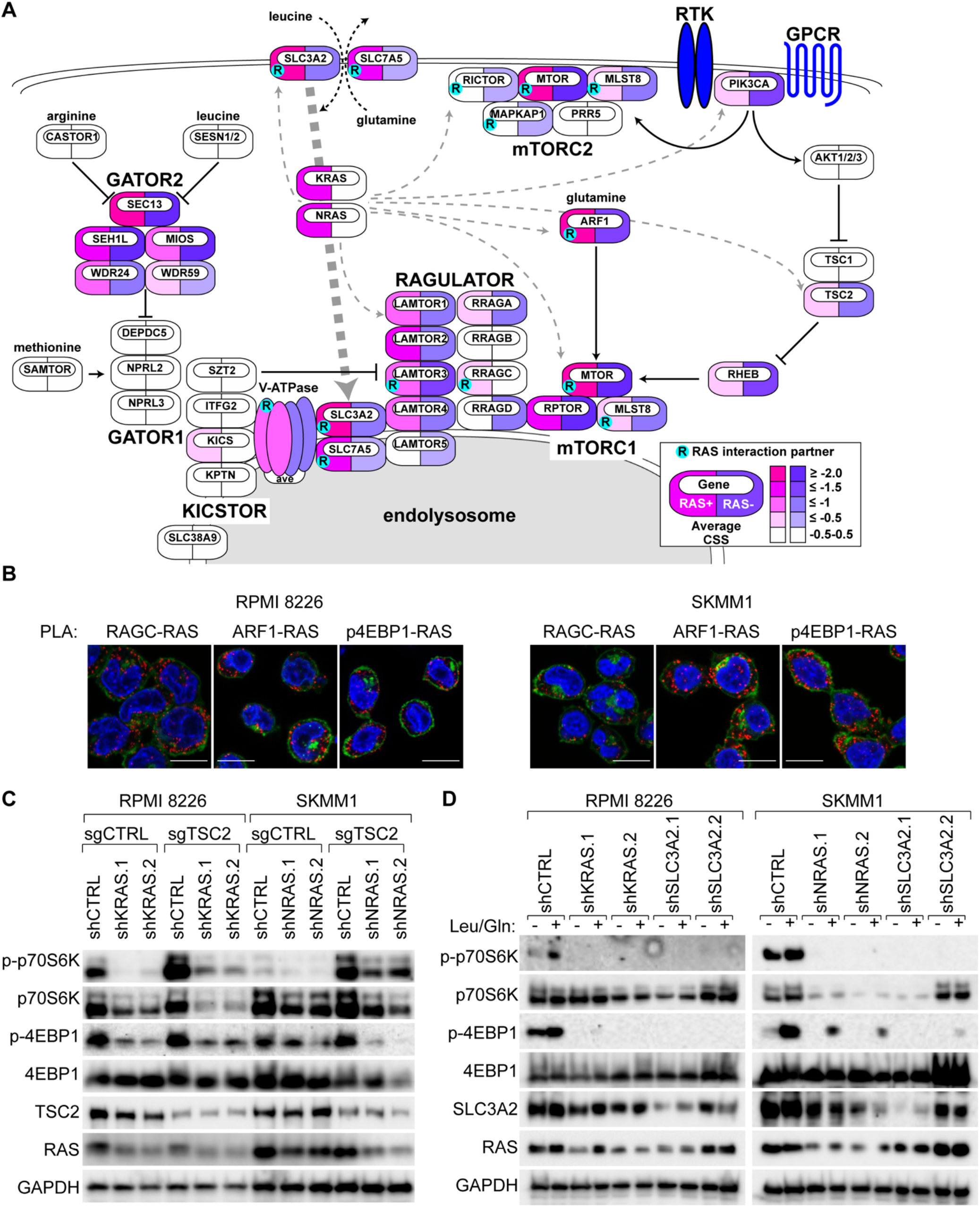
Oncogenic RAS activates mTORC1 by co-opting the amino acid sensing machinery. (A) Pathway diagram of MTOR signaling. Symbols are colored by the average CSS in RAS-dependent (pink) and RAS-independent (purple) MM cell lines, and marked with a cyan dot containing an “R” if they were found to interact with mutant KRAS and NRAS in BioID2 experiments (≥2.0 log2fc). (B) Indicated PLAs in RPMI 8226 and SKMM1 cells with PLA (red), WGA (green) and DAPI (blue). Scale bar is 10μm. Representative images from 2 or more experiments. (C) Western blot analysis of mTORC1 signaling outputs in RPMI 8226 and SKMM1 cells following KRAS or NRAS knockdown following expression of either control (sgCTRL) or TSC2 sgRNAs. Representative blot from 3 independent experiments. (D) Western blot analysis of mTORC1 signaling in RPMI 8226 (left) and SKMM1 (right) cells following amino acid starvation – or + provision of leucine and glutamine in cells expressing indicated shRNAs. Representative blot from 4 independent experiments.

Activation of mTORC1 requires concomitant engagement of nutrient sensing machinery and inhibition of TSC2, to permit RHEB association with mTORC1. TSC2 inhibition can be achieved downstream of PI3-K (*32*) or ERK (*33*). RAS has been previously described to directly bind and activate isoforms of PI3-K in solid cancers (*34*) and RAS can activate classical MAPK signaling upstream of ERK in MM (Fig 4A-C). To test if RAS is primarily regulating mTORC1 via TSC2 in MM, we examined the effect of RAS knockdown in MM cells lacking TSC2 expression. RPMI 8226 and SKMM1 cells were first transduced with control or TSC2 sgRNAs, followed by transduction with control or RAS shRNAs. We found that TSC2 deletion markedly increased phosphorylation of both 4EBP1 (S65) and p70S6K (T389), yet even these elevated levels of phosphorylation were still dependent on RAS expression (Fig. 5C), demonstrating that RAS regulates mTORC1 signaling through mechanisms other than TSC2.

Since SLC3A2-SLC7A5 is a leucine transporter and glutamine antiporter (*26*), we next tested how these amino acids regulated RAS-dependent mTORC1 activity. RPMI 8226 and SKMM1 MM cells were transduced with control shRNA or shRNAs targeting either KRAS, NRAS or SLC3A2. Following knockdown, cells were starved of amino acids for 3 hours, at which point leucine and glutamine were added back into culture or not for 90 minutes prior to lysis. Western blot analysis of mTORC1 signaling outputs found that control cells had low levels of both p70S6K (T389) and 4EBP1 (S65) phosphorylation, which was markedly enhanced by provision of leucine and glutamine (Fig. 5D). In contrast, knockdown of either RAS or SLC3A2 effectively ablated phosphorylation of p70S6K (T389) and 4EBP1 (S65) in both resting and stimulated conditions. These data demonstrate that oncogenic RAS is required for amino acid-dependent mTORC1 signaling in MM and are consistent with a model where RAS commandeers mTORC1 signaling by orchestrating components the amino acid sensing machinery.

### RAS and mTORC1 signaling in MM patients

We next sought evidence of RAS-dependent mTORC1 activity in primary MM tumors. The MTOR-RAS PLA was adapted to detect MTOR and RAS interactions in formalin-fixed paraffin-embedded (FFPE) bone marrow biopsies from a cohort of MM patients with known RAS mutation statuses. We observed numerous MTOR-RAS PLA puncta (Fig 6A, red) throughout the cytosol in CD138+ cells, a marker of plasma cells (Fig 6A, white), in a subset of MM patient samples tested. When patient samples were subdivided by RAS mutation, 33% of MM cases with KRAS or NRAS mutations (5/15; 3 KRAS, 2 NRAS) had strong MTOR-RAS PLA signals. We also found an instance where a MM case without a known RAS mutation had observable MTOR-RAS PLA (1/13; 7.7%). It is possible that this patient tumor may have represented aberrant activation of wild type RAS through other mechanism, such as overexpression and/or mutation of FGFR3, which are present ~5% of MM cases (*35*).

**Figure 6.**
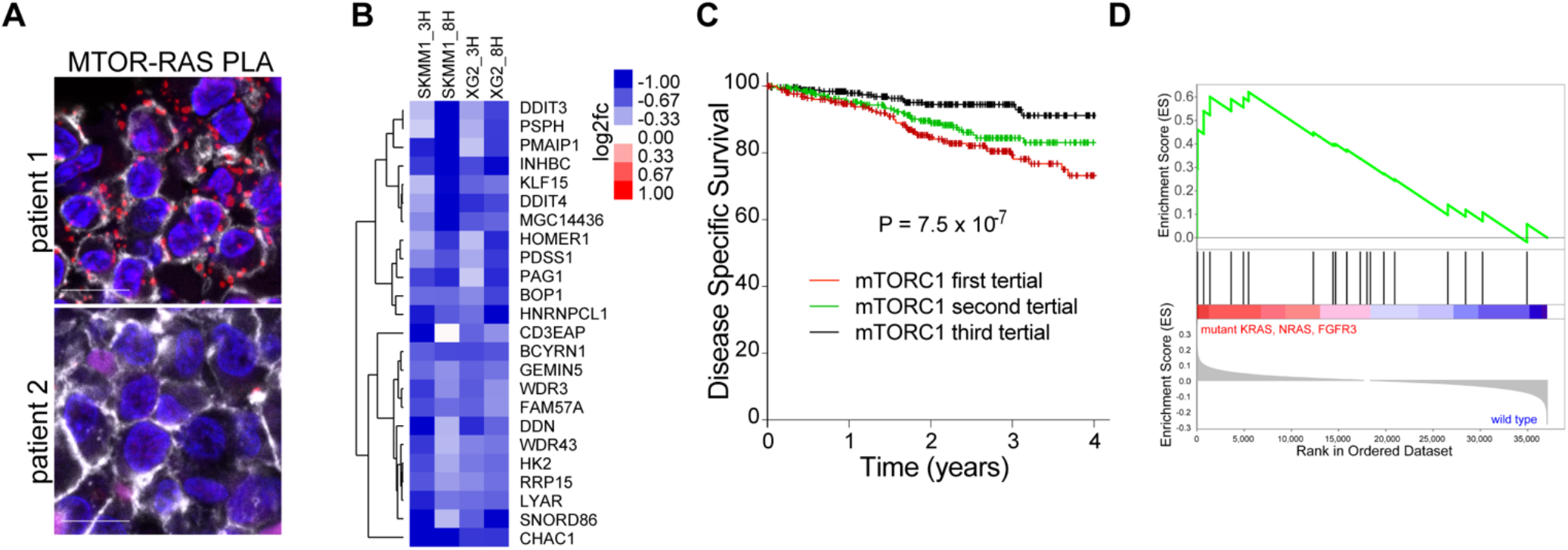
RAS-dependent mTORC1 activity in primary MM tumors. (A) PLA between MTOR and RAS in FFPE bone marrow aspirates from MM patients with PLA (red), CD138 (white) and DAPI (blue). Scale bar is 10μm. (B) Changes in expression for the mTORC1 signature genes following treatment with 100 nM everolimus for the indicated cell lines and times. (C) Kaplan-Meier survival plots of MM patients from the MMRF CoMMpass trial divided into tertials by gene expression of the mTORC1 down signature in panel B. The P-value was determined using a Cox proportional hazard model. (D) Gene Set Enrichment Analysis (GSEA) of mTORC1 signature for KRAS, NRAS and FGFR3 mutations in the MMRF CoMMpass patient cohort.

To probe for MTOR signaling in primary MM tumors, we created a gene expression signature of mTORC1-dependent genes in RAS-dependent MM cells. SKMM1 and XG2 were treated with 100 nM everolimus, and changes in gene expression relative to a DMSO control were determined at 3 and 8 hours by RNA-sequencing. Genes whose expression was decreased by an average of at least 0.5 log2fc in both cell lines were included in the signature (Fig. 6B). We applied this mTORC1 signature to gene expression data from 859 patient cases within the Multiple Myeloma Research Foundation (MMRF) CoMMpass study (*36*) and determined that it was significantly correlated with disease-specific survival in this patient cohort using a Cox proportional hazard model (p=7.5×10^−7^) (Fig. 6C), suggesting that mTORC1 signaling is correlated to poor prognosis in MM. We next performed gene set enrichment analysis (GSEA) to test if the mTORC1 signature was linked to mutations in KRAS, NRAS or FGFR3, which can activate RAS signaling independent of an oncogenic RAS mutation. Indeed, the mTORC1 signature was significantly enriched in MM samples harboring mutations in either KRAS, NRAS or FGFR3 (P=0.0058) (Fig. 6D), validating a link between oncogenic RAS signaling and mTORC1 activity in primary MM cases.

### Combined inhibition of mTORC1 and MEK1/2 is toxic to RAS-dependent MM

Our data indicated that RAS-dependent mTORC1 activity is a prominent feature in aggressive MM and would be an attractive therapeutic target to treat these cases, but mTORC1 inhibitors have had limited success as single agents in clinical trials (*37*, *38*). To improve implementation of mTORC1 inhibitors in MM, we performed a high-throughput combinatorial drug screen to evaluate synergy between everolimus and the MIPE v5.0 library of 2450 mechanistically annotated, oncology-focused compounds (*39*) in SKMM1 and RPMI 8226 cells in a series of 6×6 matrix blocks (Fig. 7A). These screens revealed exceptional synergy between everolimus and inhibitors targeting classical MAPK signaling via MEK and ERK (Fig. 7B-F; Fig. S6A-B). At the doses tested, MEK and ERK inhibitors displayed a true synthetic lethal phenotype consistent with a near *de novo* reliance on MEK and ERK signaling following inhibition of mTORC1 activity.

**Figure 7.**
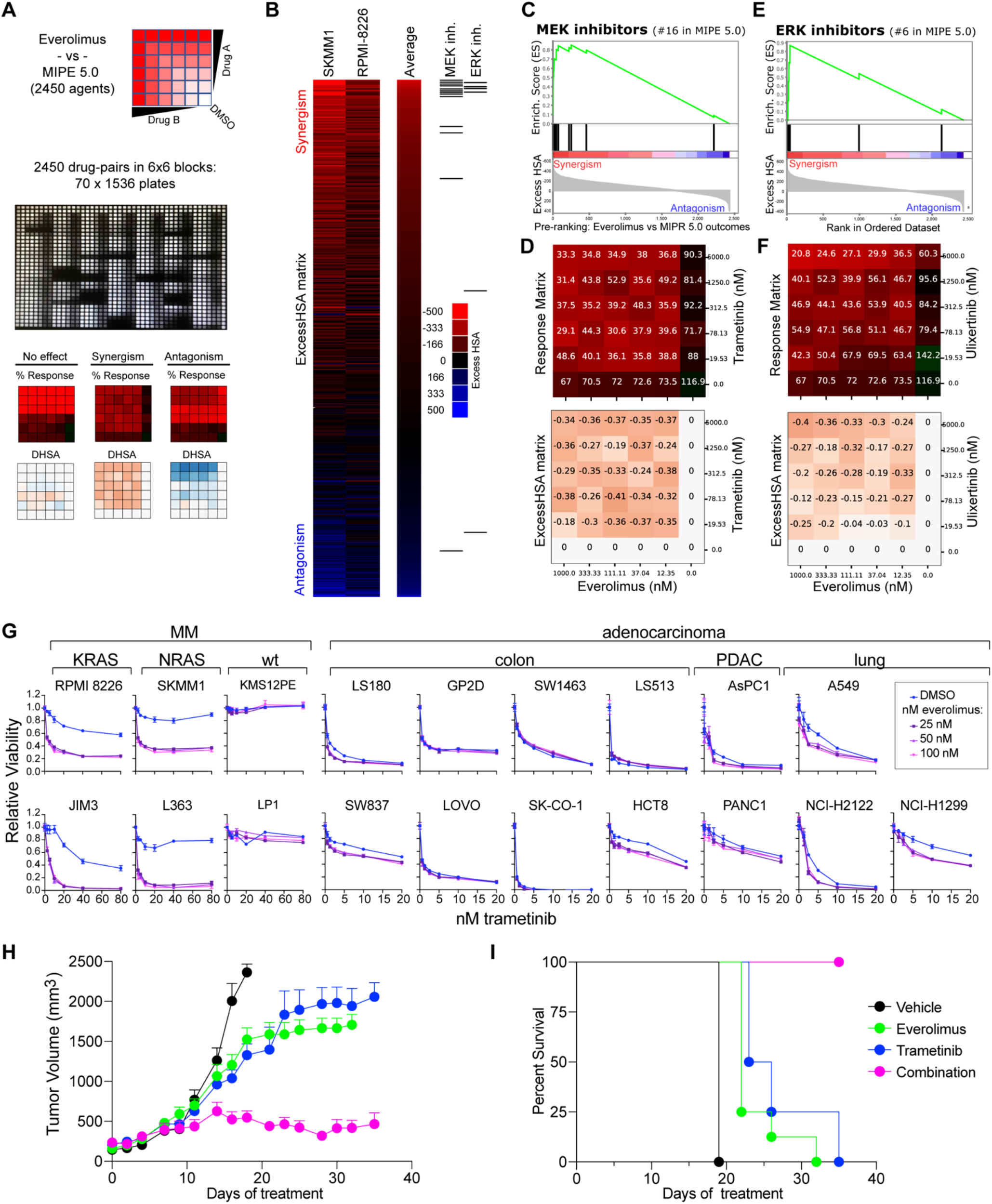
Combination therapies to target RAS-dependent mTORC1 signaling in MM. (A) Workflow for a high-throughput combinatorial drug screens comparing everolimus treatment to the MIPE 5.0 small molecule library. (B) Heat-map view of the everolimus drug-interaction landscape in SKMM1 and RPMI 8226. Drugs targeting MEK and ERK are highlighted on the right. (C) MEK inhibitors Enrichment Plot from the Drug Set Enrichment Analysis (DSEA) of the Everolimus vs MIPE5.0 screen. The average Excess HSA (SKMM1 and RPMI 8226) was used to pre-rank combinatorial outcomes before running DSEA. (D) Response matrix and ExcessHSA matrix for the everolimus vs. trametinib (MEKi) combination are shown (SKMM1). (E) ERK inhibitors Enrichment Plot from the Drug Set Enrichment Analysis (DSEA) of the everolimus vs. MIPE5.0 screen. (F) Response matrix and ExcessHSA matrix for the everolimus vs. ulixertinib (ERKi) combination are shown (SKMM1). (G) MTS viability assays for cells treated with DMSO (blue) and 25, 50 or 100nM everolimus (purples) with the indicated doses of trametinib (x-axis) for indicated MM or adenocarcinoma cell lines. (H) Tumor volume for SKMM1 xenografts treated with vehicle (black), 1 mg/kg trametinib (blue), 1mg/kg everolimus (green) or the combination (pink). Representative data from 3 independent experiments. (I) Survival for SKMM1 xenograft mice.

We evaluated combination therapy of everolimus with the MEK1/2 inhibitor, trametinib, in a cohort of MM lines (Fig. 7G, left). Consistent with the screen data, we found only modest growth inhibition from trametinib as a single agent (Fig. 7G, left, blue lines). However, combined treatment of trametinib and everolimus resulted in exceptional synergistic toxicity in all RAS-dependent MM lines tested (Fig. 7G, left, purple lines), yet we observed no drug synergy in RAS-independent MM lines (Fig. 7G). Unexpectedly, no drug synergy was detected in several adenocarcinoma cell lines harboring mutant KRAS (Fig. 7G, right), suggesting RAS-dependent activation of mTORC1 may be specific to MM. This drug combination resulted in apoptotic cell death in RAS-dependent MM lines, whereas everolimus or trametinib alone largely blocked the cell cycle (Fig. S7A), which may explain why we observe such high levels of synergistic killing with this drug combination. We conjectured that direct activation of mTORC1 by RAS and SLC3A2 is the primary mode of oncogenic RAS signaling in MM, and that RAS may not fully engage the MAPK pathway unless mTORC1 signaling is blocked. Indeed, everolimus treatment has been previously found to increase levels of phosphorylated ERK in MM (*40*), and we observed similar results (Fig. S7B).

Finally, we used mouse xenografts to determine if the combination of everolimus and trametinib retained its efficacy against MM cells *in vivo*. Combination therapy (pink) essentially halted tumor growth and was significantly more effective than either vehicle control (black), everolimus (green) or trametinib (blue) alone, without evidence of overt toxicity (Fig. 7H). In addition to inhibiting MM tumor growth, combination therapy extended survival compared to either vehicle or single agent-treated mice, and all combination mice were alive at the end of the treatment window (Fig. 7I). These data suggest that MM patients with tumors harboring active RAS signaling may specifically benefit from a combination of mTORC1 and MEK1/2 inhibitors.

## Discussion

Herein, we have described a novel mode of pathogenic RAS signaling in which RAS, SLC3A2 and MTOR comprise a signaling complex on endolysosomes that stimulates mTORC1 activity. These findings were unlocked by an unbiased proteogenomic approach that identified the essential interactomes of oncogenic RAS in MM, enabling the discovery of this unanticipated aspect of RAS biology. We propose a model in which RAS coordinates oncogenic growth and survival by subverting the amino acid sensing machinery through colocalization of SLC3A2 with MTOR and itself on endolysosomes to drive mTORC1 activity (Fig. 8). RAS-dependent activation of mTORC1 appears to be a prevalent form of pathogenic RAS signaling in MM and is distinct from RAS obliquely activating mTORC1 through activation of PI3-K (*34*). Our observations provide mechanistic insights to explain the paucity of active MEK signaling in many RAS-dependent MM tumors (*7*, *8*) and the underwhelming clinical response to MEK inhibitors in MM patients (*9*, *10*). However, we found that combinations of mTORC1 and MEK1/2 inhibitors were exceptionally synergistically toxic to RAS-dependent MM cell lines *in vitro* and nearly eliminated tumor growth in xenograft mouse models of MM. Thus, our study provides a rational basis for a combination therapy of everolimus and trametinib as an alternative to highly toxic myeloablative chemotherapy and autologous stem cell transplant in relapsed and refractory MM.

**Figure 8.**
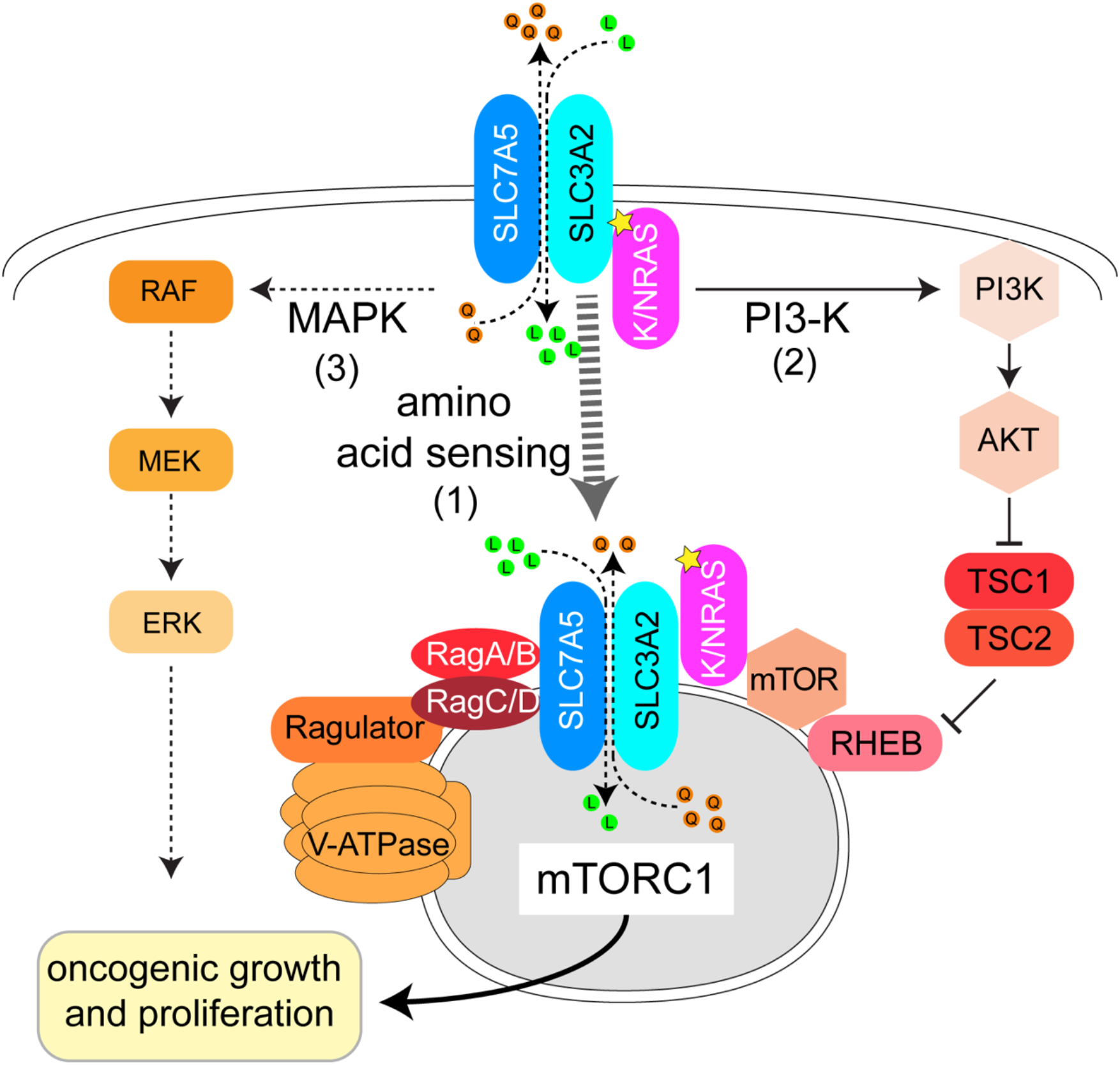
RAS co-opts mTORC1 and the amino acid sensing machinery to drive oncogenic growth in MM. Model of oncogenic RAS signaling in MM: (1) Oncogenic isoforms of RAS (KRAS or NRAS) promote localization of SLC3A2-SLC7A5 to endolysosomes with mTORC1 and RAS. (2) RAS associates with PI3-K and TSC2 to promote full activation of mTORC1 signaling. (3) RAS stimulates classical MAPK signaling.

SLC3A2 may have particular significance in MM due to the plasmacytic origin of this disease. Conditional knockout of *Slc3a2* in murine B cells resulted in a profound block in B cell proliferation and differentiation into plasma cells (*41*). Moreover, SLC3A2 and SLC7A5 expression is high in plasma cells and dependent on the transcription factor PRDM1 (Blimp-1) (*42*), a master regulator of plasma cell differentiation (*43*) and an essential MM gene (Table S1). MM patients with high levels of SLC3A2 or SLC7A5 expression by immunohistological staining had inferior progression-free survival (*25*), and a recent study found that SLC3A2 and SLC7A5 were among the most abundant proteins on the surface of MM cells (*44*). SLC3A2 may have a prominent role in plasma cells because these cells fundamentally serve as protein production factories, secreting large quantities of antibodies that require vast reserves of amino acids (*45*, *46*). Indeed, supplementation of glutamine to mice infected with *Plasmodium* led to increased numbers of long-lived plasma cells and more robust antibody responses against this pathogen (*47*). Oncogenic RAS could have taken advantage of this distinct plasma cell biology to promote tumorigenesis in MM, and expression of mutant RAS constructs in transformed lymphoblasts has been shown to be sufficient to drive plasma cell differentiation and increases antibody secretion (*48*). It remains possible that other tumor types may utilize cooperative RAS and SLC3A2. Deletion of Slc3a2 protected mice from developing tumors in a KRAS-dependent model of skin squamous cell carcinoma (*49*), and knockout of Slc7a5 prolonged survival in a KRAS-driven mouse model of colorectal cancer (*50*). However, this new mode of RAS signaling may be best exemplified in MM because of its plasma cell origins.

Targeting RAS signaling has been notoriously difficult in human cancers (*51*). Our study provides a rationale for implementing combination therapies of mTORC1 and MEK inhibitors to disrupt this unique mode of RAS signaling in MM. Similar combinations have been previously tested in clinical trials for various types of solid tumors (*52*, *53*), but most of these trials did not appreciably benefit patients (*53*). A recent study described direct interactions between RAS and components of mTORC2 in melanoma and other solid tumor cell lines (*30*). RAS-dependent mTORC2 activity would preclude the use of an mTORC1-exclusive inhibitor, and these data may explain why we observed little synergy between everolimus and trametinib in KRAS-dependent adenocarcinoma cell lines (Fig. 8B). Furthermore, these data highlight the need to consider the oncogenic cell-of-origin when designing new treatment regimens (*54*). Our data suggests that malignant cells reliant on signaling through a complex composed of RAS, SLC3A2 and MTOR are acutely sensitive to mTORC1 and MEK inhibitors. Thus, this drug combination would not be anticipated to benefit all MM patients, but would help patients with tumors that utilized cooperative RAS and mTORC1 signaling. Visualization of MTOR-RAS associations by PLA may serve as an excellent biomarker to identify patients with MM who would benefit from such combination therapies, even in the absence of known RAS mutations. In this regard, biochemical and mechanistic insights can drive the application of precision medicine strategies beyond simple mutational analysis.

## Acknowledgments

This research was supported by the Intramural Research Program of the NIH, Center for Cancer Research, National Cancer Institute. We thank Lou Staudt, Dan Hodson, Jagan Muppidi, Arthur Shaffer and Kathleen Brown for editing help and informative discussions. We also thank Hong Zhao and Weihong Xu for technical assistance.

## Author Contributions

R.M.Y conceived study; Y.Y., T.O., P.C., A.B., M.C., A.B. C.J.T., S.P. and R.M.Y. designed experiments; Y.Y. and R.M.Y created Cas9 clones and performed CRISPR screens; T.O., B.H. and R.M.Y. performed proteomic experiments; P.C. and J.Q.W. performed mouse xenografts; M.C., F.A.T., E.B., K.W., C.M., J.T., C.K.T., and C.J.T. performed or developed combinatorial drug synergy assays; G.W.W., J.D.P and D.W.H. analyzed sequencing and mutational data; G.S. processed FFPE samples; J.W.L. and C.K.V. cloned shRNAs; I.M. and D.K. obtained and processed MM clinical biopsy samples, Y.Y. and P.C. performed FACS analysis; A.B, J.W.L. and R.M.Y. performed western blot analysis or imaging studies; J.W. performed image analysis; R.M.Y wrote the manuscript; Y.Y., T.O., P.C., A.B., M.C., J.D.P., G.A.S., I.M., C.J.T., and D.K. edited the manuscript; all authors read and approved the final version of the manuscript.

## Methods

### Cell culture

All cell lines were grown at 37°C with 5% CO_2_ in advanced RPMI (Invitrogen) supplemented with fetal bovine serum (Tet tested, R&D Systems), 1% pen/strep and 1% L-glutamine (Invitrogen). Cell lines were regularly tested for mycoplasma using the MycoAlert Mycoplasma Detection Kit (Lonza) and DNA fingerprinted by examining 16 regions of copy number variants(*55*).

### Antibodies

The following antibodies were used in this study:

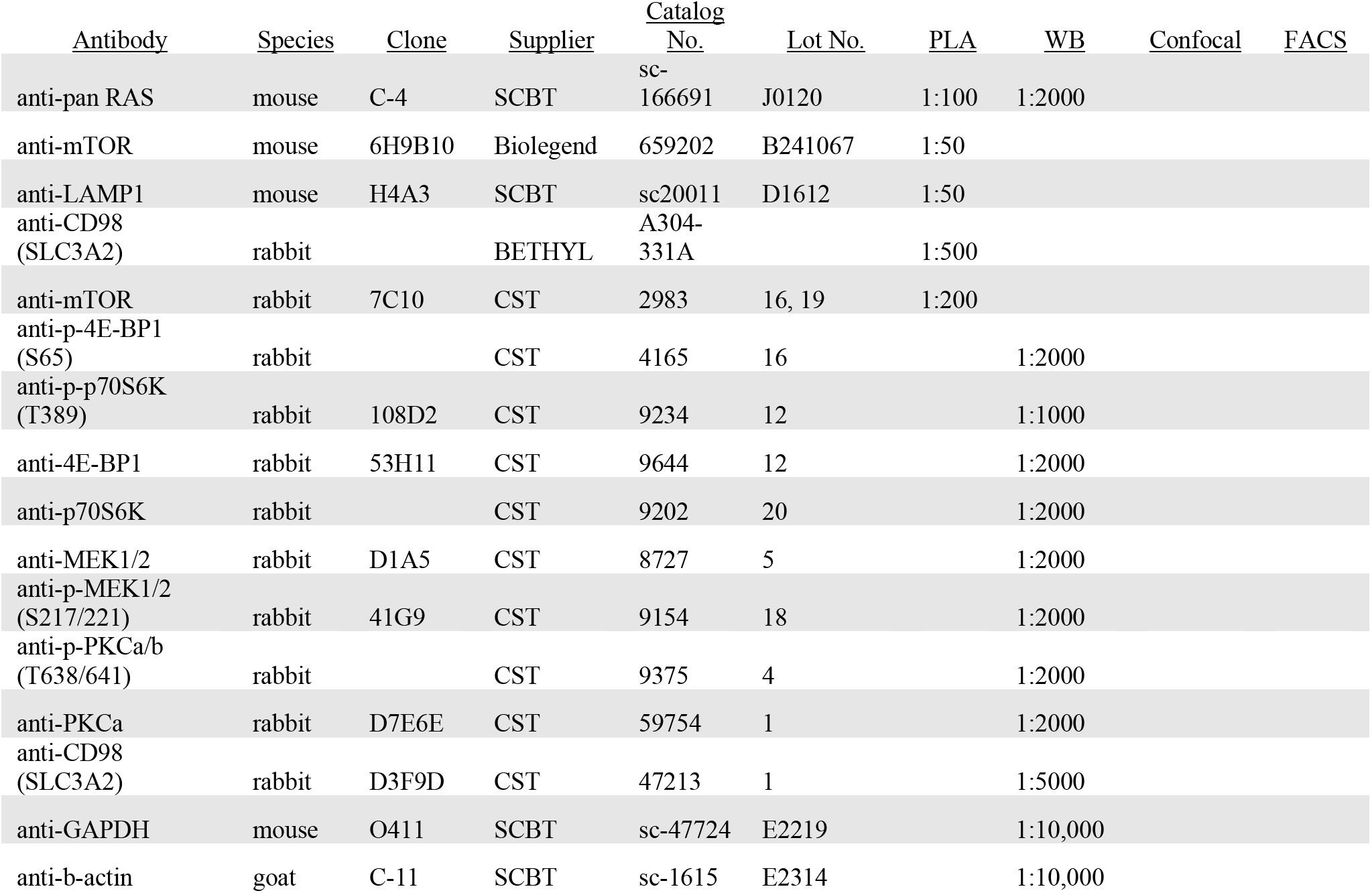
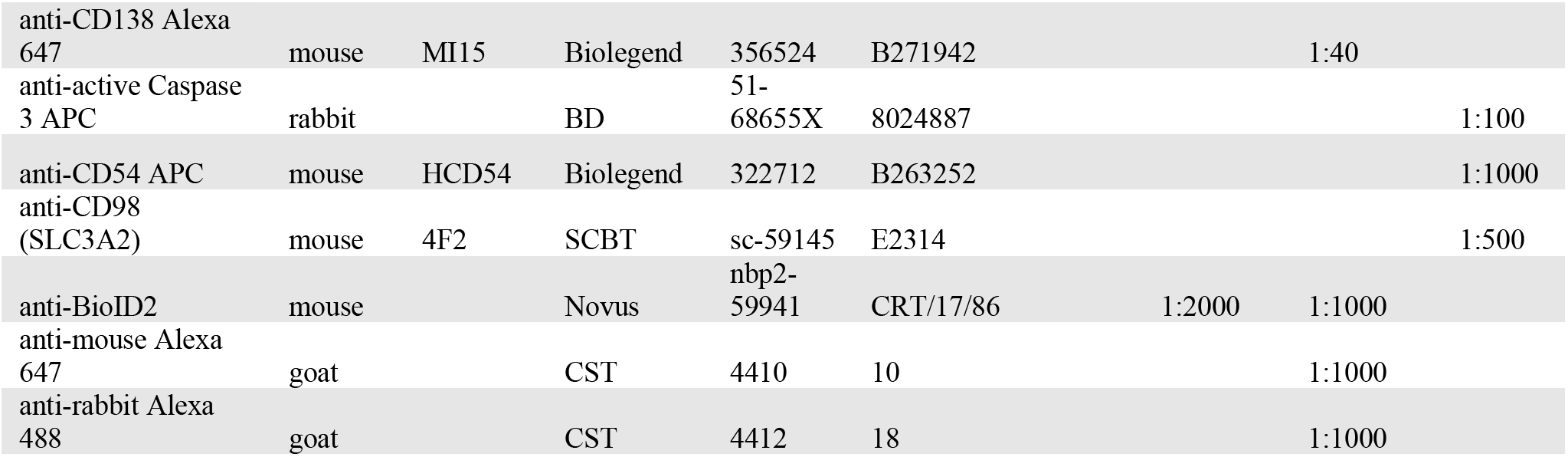

### Generation of Cas9 MM clones

MM cells were retrovirally transduced with pRetroCMV/TO-Cas9-Hygro (*13*), then selected with hygromycin and dilution cloned. Single cell clones were tested for Cas9 activity following transduction with sgRNAs for CD54 (ICAM1) or CD98 (SLC3A2) and induction with doxycycline for 10 days, at which point cells were stained for either CD54 or CD98 surface expression as determined by FACS analysis after staining with either anti-CD54 (Biolegend, clone HDCD54) or anti-CD98 (Santa Cruz Biotechnologies, clone E-5).

### CRISPR screens

CRISPR screens were performed as previously described (*13*). Briefly, lentivirus was produced from the Brunello sgRNA library (*12*) (Addgene 73178) in 293FT cells (Invitrogen) with helper plasmids pPAX2 (Addgene 12260) and pMD2.g (Addgene 12259) in Opti-Mem (Gibco) with Trans-IT 293T (Mirus). 293FT supernatants were harvested at 24, 48 and 72 hours, pre-cleared by centrifugation at 1000xg for 5 min and concentrated by 40X using Lenti-X (Takara) following the manufacturer’s instructions. Concentrated Brunello lentiviral library was added to Cas9 MM clones to yield ~ 30% infection efficiency and maintain ~1 sgRNA virus per cell with an average of 500 copies per sgRNA in total. Infected MM cells were selected with puromycin 3 days after viral transduction and allowed to grow under selection for another 3 days. At this point, 50×10^6^ cells were harvested for the day 0 timepoint and 100 ng/ml of doxycycline and 0.5 μg/ml puromycin was added to at least 50×10^6^ cells to induce Cas9 expression, after which a minimum of 50×10^6^ cells were passed every other day for 21 days to maintain an average of 500X coverage/sgRNA in the Brunello library. 50×10^6^ cells were harvested for the day 21 timepoint. DNA was extracted from Day 0 and 21 cell pellets with QIAmp DNA Blood Midi and Maxi kits (Qiagen). A nested PCR strategy was used to amplify sgRNAs from purified genomic DNA and add Next-generation sequencing adapters (Detailed methods found here:(*56*). The resulting libraries were sequenced with a NextSeq500 (Illumina), and data was processed as previously described (*13*).

### CRISPR modifier screens

SKMM1 cells were transduced with the Brunello sgRNA library as described above. Cells were selected with puromycin, and Cas9 expression was induced with doxycycline and allowed to grow for 1 week. At this point, the culture was split into 2 flasks with 50×10^6^ cells each. One flask was treated DMSO vehicle control and the other flask was treated with everolimus. Cells were treated for 2 weeks. During this time, the everolimus concentration was kept close to an IC20 dose, which ranged from 5nM at the start of treatment to 500nM after 2 weeks. After 2 weeks of treatments, 50×10^6^ cells were harvested from DMSO and everolimus treated cells, and DNA was extracted, prepared and sequenced as described above.

### CRISPR analysis

The DESeq2 algorithm (*57*) was used to estimate the log-fold change of the read count between Day 21 and Day 0 samples, or treatment and control samples, of the sgRNA guides in each cell line. Of the 77,441 guides targeting genes, 9,919 (13%) were removed for having poor performance across a large number of essential gene experiments (*13*, *58*). For each gene, the log-ratios of the remaining guides associated with that gene were averaged to estimate a gene-level, log-fold change. For each cell, these gene-level, log-fold changes were normalized by subtracting the mode of their distribution (estimated with the R-function “density”) and then divided by the root-mean-square deviation (RMSD) from that mode.

### Protein interactomes

BioID2 ((*23*); Addgene 80899) with an 8X linker of GSGGG and a SnaBI site was amplified by PCR with the following primers:

BioID2 Fwd:

~~~
AATTCGAATTCCTGAAGGGCCACCatgtatccctatgatgtgccagactatgctTTCAAGAACCTGATCTGGCTGAAGG
~~~

BioID2 Rev:

~~~
cgccggccctcgaggtacgtactaAGCGCTTCTTCTCAGGCTGAAC
~~~

The PCR fragment was purified and cloned into the StuI site in the MCS of pBMN-LYT2. The resulting BioID2-8Xlinker-pBMN-LYT2 vector permitted the addition of a BioID2-linker to the amino terminus of any gene of interest by inserting a gene at the SnaBI site using Gibson cloning (New England Biolabs). Synthetic gene fragments (G-block, IDT) of KRAS^G12V^ and NRAS^G12V^ or PCR amplicons of SLC3A2 derived from cDNA from germinal center B cells (*13*) were cloned into this vector:

KRAS^G12V^

~~~
CTGCCGGATCCGAATTCTAGCCACAatgactgaatataaacttgtggtagttggagctgTtggcgtaggcaagagtgccttg
acgatacagctaattcagaatcattttgtggacgaatatgatccaacaatagaggattcctacaggaagcaagtagtaattg
atggagaaacctgtctcttggatattctcgacacagcaggtcaagaggagtacagtgcaatgagggaccagtacatgaggac
tggggagggctttctttgtgtatttgccataaataatactaaatcatttgaagatattcaccattatagagaacaaattaaa
agagttaaggactctgaagatgtacctatggtcctagtaggaaataaatgtgatttgccttctagaacagtagacacaaaac
aggctcaggacttagcaagaagttatggaattccttttattgaaacatcagcaaagacaagacagagagtggaggatgcttt
ttatacattggtgagggagatccgacaatacagattgaaaaaaatcagcaaagaagaaaagactcctggctgtgtgaaaatt
aaaaaatgcattataatgGTAGGTGGAGGCGGGTCGGG
~~~

NRAS^G12V^

~~~
CTGCCGGATCCGAATTCTAGCCACAatgactgagtacaaactggtggtggttggagcagttggtgttgggaaaagcgcactg
acaatccagctaatccagaaccactttgtagatgaatatgatcccaccatagaggattcttacagaaaacaagtggttatag
atggtgaaacctgtttgttggacatactggatacagctggacaagaagagtacagtgccatgagagaccaatacatgaggac
aggcgaaggcttcctctgtgtatttgccatcaataatagcaagtcatttgcggatattaacctctacagggagcagattaag
cgagtaaaagactcggatgatgtacctatggtgctagtgggaaacaagtgtgatttgccaacaaggacagttgatacaaaac
aagcccacgaactggccaagagttacgggattccattcattgaaacctcagccaagaccagacagggtgttgaagatgcttt
ttacacactggtaagagaaatacgccagtaccgaatgaaaaaactcaacagcagtgatgatgggactcagggttgtatggga
ttgccatgtgtggtgatgGTAGGTGGAGGCGGGTCGGG
~~~

SLC3A2 Fwd:

~~~
CTGCCGGATCCGAATTCTAGCCACAatggagctacagcctcctgaag
~~~

SLC3A2 Rev:

~~~
CCCGACCCGCCTCCACCTACtcaggccgcgtaggggaagcg
~~~

Resultant BioID2 constructs were retrovirally transduced into MM cell lines as previously described (*13*). Transduced MM cells were purified with anti-LYT2 (mouse CD8) magnetic beads (Dynal/Thermo), and purified cells were grown in SILAC media, containing amino acids labeled with stable isotopes of arginine and lysine, for 2 weeks and then expanded to 50×10^6^ cells. In certain cases, cells were infected with pLKO-shKRAS.2 or PLKO-shNRAS.1 (see below). 16 hours prior to lysis, biotin (Sigma) was added to a final concentration of 50μM to transduced cells. Cells were then lysed at 1 × 10^7^ cells per ml in RIPA buffer modified for MS analysis (1% NP-40, 0.5% deoxycholate, 50 mM Tris, pH 7.5, 150 mM NaCl, 1 mM Na_3_VO_4_, 5mM NaF, 1 mM AEBSF) for 10 min. on ice. Lysates were cleared by centrifugation at 14,000xg for 20 min. at 4°C. 35μl of pre-washed streptavidin agarose beads (Thermo) were added to each sample; samples were then rotated at 4°C for 2 hours, then washed three times in 1X RIPA buffer, then solubilized with 4X LDS sample buffer (Invitrogen) with 1% Nupage reducing agent (Invitrogen), and boiled for 5 min.

For MS analysis, proteins were separated by one-dimensional gel electrophoresis (4–12% NuPAGE Bis-Tris Gel; Invitrogen), and the entire lane of a Coomassie blue-stained gel was cut into 23 slices. All slices were processed as described previously (*59*). After tryptic digestion of the proteins the resulting peptides were resuspended in sample loading buffer (2% acetonitrile and 0.05% trifluoroacetic acid) and were separated by an UltiMate 3000 RSLCnano HPLC system (Thermo Fisher Scientific) coupled online to a Q Exactive HF mass spectrometer (Thermo Fisher Scientific). First, peptides were desalted on a reverse phase C18 precolumn (Dionex 5 mm length, 0.3 mm inner diameter) for 3 minutes. After 3 minutes the precolumn was switched online to the analytical column (30cm length, 75 mm inner diameter) prepared in-house using ReproSil-Pur C18 AQ 1.9 mm reversed phase resin (Dr. Maisch GmbH). Buffer A consisted of 0.1 % formic acid in H_2_O, and buffer B consisted of 80% acetonitrile and 0.1% formic acid in H_2_O. The peptides eluted from buffer B (5 to 42 % gradient) at a flow rate of 300 nl/min over 76 min. The temperature of the precolumn and the analytical column was set to 50°C during the chromatography. The mass spectrometer was operated in a TopN data-dependent mode, where the 30 most intense precursors from survey MS1 scans were selected with an isolation window of 1.6 Th for MS2 fragmentation under a normalized collision energy of 28. Only precursor ions with a charge state between 2 and 5 were selected. MS1 scans were acquired with a mass range from 350 to 1600 m/z at a resolution of 60,000 at 200 m/z. MS2 scans were acquired with a starting mass of 110 Th at a resolution of 15,000 at 200 m/z with maximum IT of 54ms. AGC targets for MS1 and MS2 scans were set to 1E6 and 1E5, respectively. Dynamic exclusion was set to 20 seconds.

### mNeonGreen fusions

For co-immunoprecipitation studies, mutant RAS isoforms of KRAS^G12D^, NRAS^G12D^ and NRAS^L61Q^ were linked to mNeonGreen on their N-terminus were cloned into pBMN at the StuI sites. MM cells were retrovirally transduced and selected for LYT2 expression, as described above. For lysis, 20 × 10^6^ cells were lysed in 0.5% CHAPS lysis buffer (50 mM Tris, pH 7.5, 150 mM NaCl, 1 mM Na_3_VO_4_, 5mM NaF, 1 mM AEBSF) for 10 min. on ice, and lysates were cleared by centrifugation at 14,000xg for 20 min. at 4°C and the post-nuclear supernatant was collected. Samples were divided in two and 25 μl of mNeonGreen-Trap agarose (Chromotek) was added to pulldown mNeonGreen-tagged RAS constructs, or 25 μl of saturated control beads (Chromotek), after which lysates were rotated at 4C for 2 hours. Beads were then washed 3X in CHAPS lysis buffer and 30 μl 2X Laemmli sample buffer (BioRad) was added to each sample, followed by boiling for 5 min. Samples were then subjected to western blot analysis as described below.

Wild type and mutant versions of SLC3A2 were cloned in a similar fashion and retrovirally expressed in MM cells.

mNeonGreen-SLC3A2 wild type

~~~
atggagctacagcctcctgaagcctcgatcgccgtcgtgtcgattccgcgccagttgcctggctcacattcggaggctggtg
tccagggtctcagcgcgggggacgactcagagacggggtctgactgtgttacccaggctggtcttcaactcttggcctcaag
tgatcctcctgccttagcttccaagaatgctgaggttacagtagaaacggggtttcaccatgttagccaggctgatattgaa
ttcctgacctcaattgatccgactgcctcggcctccggaagtgctgggattacaggcaccatgagccaggacaccgaggtgg
atatgaaggaggtggagctgaatgagttagagcccgagaagcagccgatgaacgcggcgtctggggcggccatgtccctggc
gggagccgagaagaatggtctggtgaagatcaaggtggcggaagacgaggcggaggcggcagccgcggctaagttcacgggc
ctgtccaaggaggagctgctgaaggtggcaggcagccccggctgggtacgcacccgctgggcactgctgctgctcttctggc
tcggctggctcggcatgcttgctggtgccgtggtcataatcgtgcgagcgccgcgttgtcgcgagctaccggcgcagaagtg
gtggcacacgggcgccctctaccgcatcggcgaccttcaggccttccagggccacggcgcgggcaacctggcgggtctgaag
gggcgtctcgattacctgagctctctgaaggtgaagggccttgtgctgggtccaattcacaagaaccagaaggatgatgtcg
ctcagactgacttgctgcagatcgaccccaattttggctccaaggaagattttgacagtctcttgcaatcggctaaaaaaaa
gagcatccgtgtcattctggaccttactcccaactaccggggtgagaactcgtggttctccactcaggttgacactgtggcc
accaaggtgaaggatgctctggagttttggctgcaagctggcgtggatgggttccaggttcgggacatagagaatctgaagg
atgcatcctcattcttggctgagtggcaaaatatcaccaagggcttcagtgaagacaggctcttgattgcggggactaactc
ctccgaccttcagcagatcctgagcctactcgaatccaacaaagacttgctgttgactagctcatacctgtctgattctggt
tctactggggagcatacaaaatccctagtcacacagtatttgaatgccactggcaatcgctggtgcagctggagtttgtctc
aggcaaggctcctgacttccttcttgccggctcaacttctccgactctaccagctgatgctcttcaccctgccagggacccc
tgttttcagctacggggatgagattggcctggatgcagctgcccttcctggacagcctatggaggctccagtcatgctgtgg
gatgagtccagcttccctgacatcccaggggctgtaagtgccaacatgactgtgaagggccagagtgaagaccctggctccc
tcctttccttgttccggcggctgagtgaccagcggagtaaggagcgctccctactgcatggggacttccacgcgttctccgc
tgggcctggactcttctcctatatccgccactgggaccagaatgagcgttttctggtagtgcttaactttggggatgtgggc
ctctcggctggactgcaggcctccgacctgcctgccagcgccagcctgccagccaaggctgacctcctgctcagcacccagc
caggccgtgaggagggctcccctcttgagctggaacgcctgaaactggagcctcacgaagggctgctgctccgcttccccta
cgcggcctga
~~~

mNeonGreen-SLC3A2
K532E

~~~
atggagctacagcctcctgaagcctcgatcgccgtcgtgtcgattccgcgccagttgcctggctcacattcggaggctggtg
tccagggtctcagcgcgggggacgactcagagacggggtctgactgtgttacccaggctggtcttcaactcttggcctcaag
tgatcctcctgccttagcttccaagaatgctgaggttacagtagaaacggggtttcaccatgttagccaggctgatattgaa
ttcctgacctcaattgatccgactgcctcggcctccggaagtgctgggattacaggcaccatgagccaggacaccgaggtgg
atatgaaggaggtggagctgaatgagttagagcccgagaagcagccgatgaacgcggcgtctggggcggccatgtccctggc
gggagccgagaagaatggtctggtgaagatcaaggtggcggaagacgaggcggaggcggcagccgcggctaagttcacgggc
ctgtccaaggaggagctgctgaaggtggcaggcagccccggctgggtacgcacccgctgggcactgctgctgctcttctggc
tcggctggctcggcatgcttgctggtgccgtggtcataatcgtgcgagcgccgcgttgtcgcgagctaccggcgcagaagtg
gtggcacacgggcgccctctaccgcatcggcgaccttcaggccttccagggccacggcgcgggcaacctggcgggtctgaag
gggcgtctcgattacctgagctctctgaaggtgaagggccttgtgctgggtccaattcacaagaaccagaaggatgatgtcg
ctcagactgacttgctgcagatcgaccccaattttggctccaaggaagattttgacagtctcttgcaatcggctaaaaaaaa
gagcatccgtgtcattctggaccttactcccaactaccggggtgagaactcgtggttctccactcaggttgacactgtggcc
accaaggtgaaggatgctctggagttttggctgcaagctggcgtggatgggttccaggttcgggacatagagaatctgaagg
atgcatcctcattcttggctgagtggcaaaatatcaccaagggcttcagtgaagacaggctcttgattgcggggactaactc
ctccgaccttcagcagatcctgagcctactcgaatccaacaaagacttgctgttgactagctcatacctgtctgattctggt
tctactggggagcatacaaaatccctagtcacacagtatttgaatgccactggcaatcgctggtgcagctggagtttgtctc
aggcaaggctcctgacttccttcttgccggctcaacttctccgactctaccagctgatgctcttcaccctgccagggacccc
tgttttcagctacggggatgagattggcctggatgcagctgcccttcctggacagcctatggaggctccagtcatgctgtgg
gatgagtccagcttccctgacatcccaggggctgtaagtgccaacatgactgtgaagggccagagtgaagaccctggctccc
tcctttccttgttccggcggctgagtgaccagcggagtgaggagcgctccctactgcatggggacttccacgcgttctccgc
tgggcctggactcttctcctatatccgccactgggaccagaatgagcgttttctggtagtgcttaactttggggatgtgggc
ctctcggctggactgcaggcctccgacctgcctgccagcgccagcctgccagccaaggctgacctcctgctcagcacccagc
caggccgtgaggagggctcccctcttgagctggaacgcctgaaactggagcctcacgaagggctgctgctccgcttccccta
cgcggcctga
~~~

mNeonGreen-SLC3A2 truncation

~~~
atgcccggctgggtacgcacccgctgggcactgctgctgctcttctggctcggctggctcggcatgcttgctggtgccgtgg
tcataatcgtgcgagcgccgcgttgtcgcgagctaccggcgcagaagtggtggcacacgggcgccctctaccgcatcggcga
ccttcaggccttccagggccacggcgcgggcaacctggcgggtctgaaggggcgtctcgattacctgagctctctgaaggtg
aagggccttgtgctgggtccaattcacaagaaccagaaggatgatgtcgctcagactgacttgctgcagatcgaccccaatt
ttggctccaaggaagattttgacagtctcttgcaatcggctaaaaaaaagagcatccgtgtcattctggaccttactcccaa
ctaccggggtgagaactcgtggttctccactcaggttgacactgtggccaccaaggtgaaggatgctctggagttttggctg
caagctggcgtggatgggttccaggttcgggacatagagaatctgaaggatgcatcctcattcttggctgagtggcaaaata
tcaccaagggcttcagtgaagacaggctcttgattgcggggactaactcctccgaccttcagcagatcctgagcctactcga
atccaacaaagacttgctgttgactagctcatacctgtctgattctggttctactggggagcatacaaaatccctagtcaca
cagtatttgaatgccactggcaatcgctggtgcagctggagtttgtctcaggcaaggctcctgacttccttcttgccggctc
aacttctccgactctaccagctgatgctcttcaccctgccagggacccctgttttcagctacggggatgagattggcctgga
tgcagctgcccttcctggacagcctatggaggctccagtcatgctgtgggatgagtccagcttccctgacatcccaggggct
gtaagtgccaacatgactgtgaagggccagagtgaagaccctggctccctcctttccttgttccggcggctgagtgaccagc
ggagtaaggagcgctccctactgcatggggacttccacgcgttctccgctgggcctggactcttctcctatatccgccactg
ggaccagaatgagcgttttctggtagtgcttaactttggggatgtgggcctctcggctggactgcaggcctccgacctgcct
gccagcgccagcctgccagccaaggctgacctcctgctcagcacccagccaggccgtgaggagggctcccctcttgagctgg
aacgcctgaaactggagcctcacgaagggctgctgctccgcttcccctacgcggcctga
~~~

### MS data analysis

MS data analysis was performed using the software MaxQuant (version 1.6.0.1) linked to the UniProtKB/Swiss-Prot human database containing 155990 protein entries and supplemented with 245 frequently observed contaminants via the Andromeda search engine.(*60*) Precursor and fragment ion mass tolerances were set to 6 and 20 ppm after initial recalibration, respectively. Protein biotinylation, N-terminal acetylation and methionine oxidation were allowed as variable modifications. Cysteine carbamidomethylation was defined as a fixed modification. Minimal peptide length was set to 7 amino acids, with a maximum of two missed cleavages. The false discovery rate (FDR) was set to 1% on both the peptide and the protein level using a forward-and-reverse concatenated decoy database approach. For SILAC quantification, multiplicity was set to two or three for double (Lys+0/Arg+0, Lys+8/Arg+10) or triple (Lys+0/Arg+0, Lys+4/Arg+6, Lys+8/Arg+10) labeling, respectively. At least two ratio counts were required for peptide quantification. The “re-quantify” option of MaxQuant was enabled. Data was filtered for low confidence peptides.

### Phosphoproteome Analysis

SKMM1 cells were grown and expanded in SILAC media to 100 × 10^6^ cells per condition. At this point, cells were transduced with concentrated lentivirus: cells in ‘Light’ media were transduced with shCtrl, ‘Medium’ were transduced with shNRAS.1 and ‘Heavy’ were transduced with shNRAS.2 (See below). Cells were selected with puromycin (Invitrogen) the following day and allowed to grow under selection conditions for 2 days, after which they were lysed in 1% NP-4050 mM Tris, pH 7.5, 150 mM NaCl, 1 mM Na_3_VO_4_, 5mM NaF with 1 tablet/10 ml EDTA-free protease inhibitor cocktail tablets (Roche). Changes in global phosphorylation were analyzed as previously described (*61*). For analysis, the log2-fold change of shNRAS.1 and shNRAS.2 were averaged.

### Pathway Analysis

Pathway analysis was performed using ToppFun from the ToppGene Suite (*62*). Gene Ontogeny Biological Process and Pathway analysis were used with the indicated gene lists and log2fc values. Additional analysis was performed using Metacore+MetaDrug™ version 21.3. by applying the integrated Enrichment Analysis Workflow, Pathway Maps, Process Networks, Metabolic Networks and GO Processes of identified RAS interactors (cut-off: log2fc +1.5).

### shRNA and sgRNA mediated knockdown

Individual shRNAs were obtained from the MISSION shRNA Library from the RNAi Consortium TRC1.0 in the pLKO.1 vector (SIGMA):

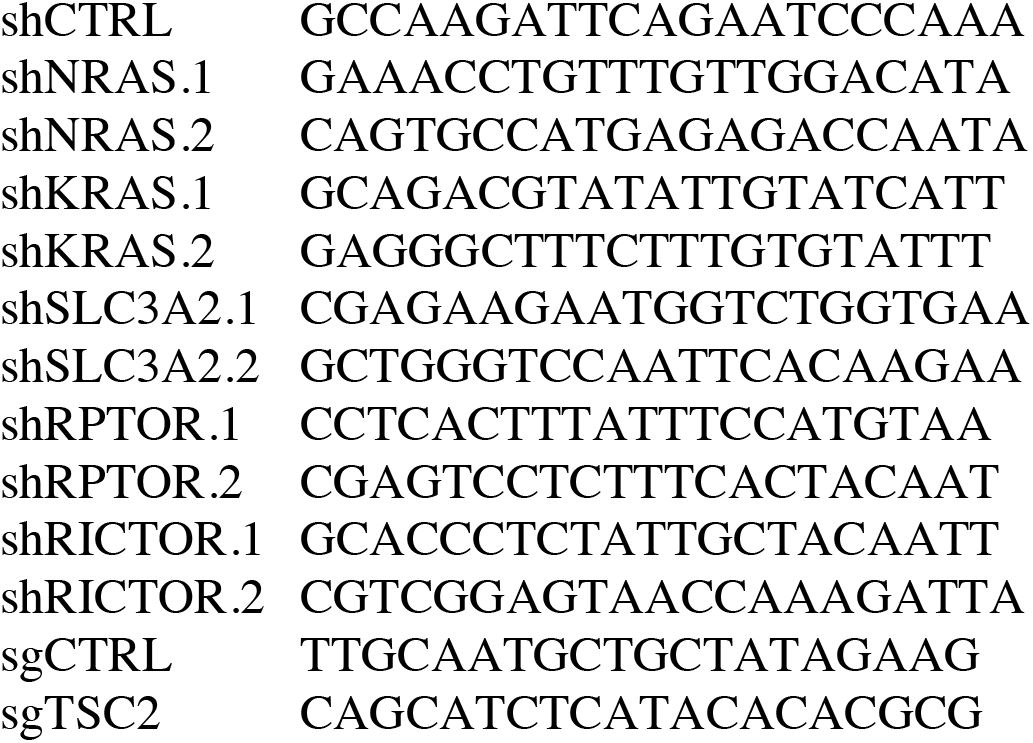

Lentiviral transductions of shRNAs were performed as described above. Transduced MM cells were selected with 1μg/ml puromycin (Gibco) for 2 days before either western blot analysis or proximity ligation.

### Proximity Ligation Assay

MM cells were transduced with shRNAs or treated with 50nM everolimus (Sellekchem) or 5μM JPH203 (SIGMA) were plated onto a 15 well μ-Slide Angiogenesis ibiTreat chamber slide (Ibidi) and allowed to adhere to the surface for 1 hour at 37°C. Cells were next fixed with 4% paraformaldehyde (Electron Microscopy Sciences) for 20 min at room temperature and then washed in PBS (Invitrogen). Cellular membranes were labeled with 5 μg/ml wheat germ agglutinin (WGA) conjugated to Alexa Fluor 488 (Thermo Fisher Scientific) for 10 min at room temperature. Cells were permeabilized in cold methanol for 20 min, washed in PBS and then blocked in Duolink Blocking buffer (Sigma) for 1 hour at room temperature. Primary antibodies were diluted in Duolink Antibody Diluent (Sigma) and incubated overnight at 4°C (See above). The following morning, cells were washed for 20 min in TBST with 0.5% tween-20, followed by addition of the appropriate Duolink secondary antibodies (Sigma), diluted and mixed according to the manufacturer’s instructions. Cells were incubated for 1 hour at 37°C, after which cells were washed in TBST with 0.5% tween-20 for 10 min. For studies examining immunofluorescence and PLA simultaneously, we incubated samples with labeled antimouse and anti-rabbit secondary antibodies for 1 hour at room temperature, followed by 10 min. wash in TBST with 0.5% tween-20. Ligation and amplification steps of the PLA were performed using the Duolink in situ Detection Reagents Orange kit (Sigma) according to the manufacturer’s instructions. Following the PLA, cells were mounted in Fluoroshield Mounting Medium with DAPI (Abcam). Images were acquired on a Zeiss LSM 880 Confocal microscope using Zeiss Zen Black version 2.3. Images for display were prepared with NIH ImageJ/FIJI software version 2.0.0-rc-65/1.5ls (*63*). PLA spots were counted using Blobfinder version 3.2 (*64*). PLA Score was determined by normalizing the number of PLA spots counted in each sample to the average number of PLA spots counted in the control sample, which was set to 100. Box and whisker plots display the median PLA Score with whiskers incorporating 5-95% of all data; outliers are displayed as dots. Statistical comparisons were made by one-way ANOVA using Prism 9.

The PLA was performed on formalin-fixed, paraffin-embedded (FFPE) tissue microarrays or biopsy samples in a similar manner. Samples were deparaffinized in xylene and rehydrated in graded alcohol and distilled water. Heat induced antigen retrieval was performed at pH 6.0 for 30 minutes. Slides were then placed in tris-buffered solution and prepared for proximity ligation assay, as described above, MM samples were co-stained with mouse anti-human CD138-Alexa647 (Biolegend, clone MI15). PLA was scored manually in CD138+ cells in a blinded fashion as either – or +. All primary patient samples were anonymized or de-identified for subsequent PLA analysis.

### Human Samples

All cases were either needle aspirates from bone marrow or bone marrow aspirate clots. Samples were fixed in 10% buffered formalin for 18-24 hours and paraffin embedded for long term storage. Samples were studied in accordance with the ethics and principles of the Declaration of Helsinki and under Institutional Review Board approved protocols from the National Cancer Institute National Institutes of Health Protocol Review Office (protocol number 11-C-0221). All samples were anonymized or deidentified for subsequent PLA analysis.

### Western blot analysis

Cells were then lysed at 1 × 10^7^ cells per ml in modified RIPA buffer (1% NP-40, 0.5% deoxycholate, 50 mM Tris, pH 7.5, 150 mM NaCl, 1 mM Na_3_VO_4_, 5mM NaF, 1 mM AEBSF) for 10 min. on ice. Lysates were cleared by centrifugation at 14,000xg for 20 min. at 4°C, and the post-nuclear supernatant was collected. Protein concentrations were determined using the Pierce BCA protein assay kit (Thermo) according to the manufacturers protocol. 100 ul of lysate and 40 μl of 4X Laemmli sample buffer (BioRad) with 1% β-mercaptoethanol (BioRad) were combined and then boiled for 5 min. 15 μg of eacg lysate was run on a 4-12% gradient gel (BioRad) and transferred to a PVDF membrane (Millipore) on an Owl semi-dry transfer device (Thermo). PVDF membranes were blocked with 5% milk (BioRad) in TBST and then probed with listed antibodies diluted in either 1% BSA (anti-phospho-specific antibodies; MPI) or milk and anti-rabbit-HRP or anti-mouse-HRP (Cell Signaling Technology) where appropriate.

### Amino acid and SDF1 stimulations

For amino acid stimulations, RPMI 8226 and SKMM1 cells were transduced with shRNAs as described above. Two days after puromycin selection, cells were washed twice with PBS to remove growth media and cells were re-plated with Tyrode’s buffer (120 mM NaCl, 5 mM KCl, 25 mM HEPES, 2 mM CaCL2, 2 mM MgCL2, 6 g/L glucose, pH 7.4) with 4% dialyzed FBS (Sigma). Cells were grown at 37°C for 3 hours under these conditions. At this point, 10^6^ cells were placed in 1 ml Tyrode’s:FBS, and cells were either left unstimulated or provided 10 μl of 100X leucine/glutamine (final concentration: L-leucine 50mg/L (Sigma); L-glutamine 300mg/L (Sigma)). Cells were then incubated at 37°C for 90 minutes, at which point there were lysed in SDS sample buffer and subjected to western blot analysis as described above. For drug experiments, all drugs were obtained from Selleckchem and used at the indicated concentrations.

For SDF1 stimulation, 10^6^ cells were placed in 1 ml of advanced RPMI with 4% FBS at 37°C. Cells were either stimulated with 100 ng/ml SDF1 (Peprotech) or an equivalent volume of PBS for 5 min. Then, cells were lysed in SDS sample buffer and subjected to western blot analysis as described above.

### FACS analysis

MM cell lines were transduced with sgCD54 vector co-expressing GFP and stained with anti-CD54 (1:1000) to select Cas9-expressing cells as previously described (*13*). SLC3A2 surface expression was measured by staining 2×10^5^ cells on ice with anti-CD98 (1:500) for 20 minutes in FACS buffer (PBS with 2% BSA). Cells were washed with FACS buffer and stained with anti-mouse-Alexa647 (1:1000; CST) for 20 minutes on ice, then washed again and resuspended in 250 μl of FACS buffer. These cells were analyzed on a BD FACS Calibur using CellQuest Pro version 6.0 and analyzed with FlowJo version 10. For cell cycle analysis, cells were treated for 1 day with DMSO, 50 nM everolimus (Selleckchem), 5 nM trametinib (Selleckchem) or both drugs together. Treated cells were stained with DyeCycle Violet450 (Invitrogen) following the manufacturer’s protocol. For cell viability analysis, cells were treated for 2 days under the same conditions and then stained with either 7AAD (Invitrogen) and Annexin V-PE (BD) or stained with anti-cleaved caspase 3 Alexa 647 (BD), following the manufacturer’s protocol. Stained cells were analyzed with a CytoFLEX LX (Beckman Coulter) and data was analyzed with FlowJo version 10.

### Drug Sensitivity Assays

MM cell lines were seeded at ~5000 cells/well in triplicate in 96-well plates. Trametinib and everolimus (SelleckChem) dissolved in DMSO were diluted in equal volumes at the indicated concentrations. Cells were cultured with drugs for 4 days. Drugs were replenished after 48 hrs. Metabolic activity was measured at day 4 with CellTiter 96 (Promega) following the manufacturer’s protocol. Absorbance was measured at 490nm using a 96-well Tecan Infinite 200 Pro plate reader.

### Gene Expression Profiling and Signature Enrichment

SKMM1 and XG2 MM cells were treated with 50 nM everolimus and harvested at indicated times after shRNA induction. RNA was isolated using AllPrep kits (Qiagen) and sequencing libraries were prepared using the TruSeq RNA Library Prep Kit V2 (Illumina). Sequencing and analysis to obtain digital gene expression were previously described (*58*). Changes in gene expression between everolimus treated and DMSO control cells were determined, and genes with an average log2 fold change of less than −0.5 per in both cell lines were included in the mTORC1 signature. Digital gene expression (DGE) values and gene-signature averages were calculated as previously described (*58*). P-values for differences in signature averages were calculated using a two-sided t-test. P-values for the association between mTORC1 and survival were from a two-sided likelihood-ratio test based on a Cox proportional hazard model with the mTORC1 signature treated as a continuous variable.

### CoMMpass Data

Data from the MMRF CoMMpass dataset (*36*) was downloaded through GDC portal using GDC-client tool and processed the GDC standard pipelines (https://docs.gdc.cancer.gov/Data/Bioinformatics_Pipelines/DNA_Seq_Variant_Calling_Pipeline/). Then, processed WES result was further annotated as previously described (*58*) to call gene mutations. RNA-Seq was further processed and normalized as previously described (*58*).

### Quantitative high-throughput combination screening (qHTCS)

Drug combination screening was performed as previously described (*65*). Briefly, 10 nL of compounds were acoustically dispensed into 1536-well white polystyrene tissue culture-treated plates with an Echo 550 acoustic liquid handler (Labcyte). Cells were then added to compound-containing plates at a density of 500-cells/well in 5 μL of medium. A 5-point custom concentration range, with constant 1:4 dilution was used for all the MIPE 5.0 drugs (*39*) in the primary 6×6 matrix screening against Everolimus (1:3 dilution), and a 9-point custom concentration range was used for secondary validation in 10×10 matrix format.

Plates were incubated for 48 hours at standard incubator conditions covered by a stainless steel gasketed lid to prevent evaporation. 48h post compound addition, 3 μL of Cell Titer Glo (Promega) were added to each well, and plates were incubated at room temperature for 15 minutes with the stainless-steel lid in place. Luminescence readings were taken using a Viewlux imager (PerkinElmer) with a 2 second exposure time per plate.

### Xenograft

All mouse experiments were approved by the National Cancer Institute Animal Care and Use Committee (NCI-ACUC) and were performed in accordance with NCI-ACUC guidelines and under approved protocols. Female NSG (non-obese diabetic/severe combined immunodeficient/common gamma chain deficient) mice were obtained from NCI Fredrick Biological Testing Branch and used for the xenograft experiments between 6–8 weeks of age. SKMM1 and JIM3 multiple myeloma tumors were established by subcutaneous injection of 10 × 10^6^ cells in a 1:1 Matrigel/PBS suspension. Treatments were initiated when tumor volume reached a mean of 200mm^3^. MEK inhibitor (trametinib; Selleckchem) was prepared in 10% (v/v) DMSO + 90% (v/v) corn oil and administered p.o. once per day (1mg/kg/day). mTORC1 inhibitor (everolimus; Selleckchem) was prepared in 10% (v/v) DMSO + 30% (v/v) propylene glycol + 5% (v/v) Tween 80 + 55% (v/v) H2O and administered p.o. once per day (1mg/kg/day). For the MEK/MTOR inhibitor combination, drugs were given at the same concentration and schedule as single agents. Tumor growth was monitored every other day by measuring tumor size in two orthogonal dimensions and tumor volume was calculated by the following equation: tumor volume = (length × width2)/2.

### Data availability

All CRISPR, proteomics and gene expression datasets are available in the supplementary tables.

## Notes

The authors declare no potential conflicts of interest.

### Competing Interest Statement

The authors have declared no competing interest.

### Summary of Updates

Revised text and updated data in figures 5 and S3.

