## Supplemental Figures and Legends for "Oncogenic RAS commandeers amino acid sensing machinery to aberrantly activate mTORC1 in multiple myeloma"

**Supplemental Figure Legends**

**Supplemental Figure 1. CRISPR screens identify essential and tumor suppressor genes in multiple myeloma.** (A) CSS (y-axis) for indicated genes. Each dot is an individual MM cell line from the cohort of 17 cell lines identified in Fig. 1A. (B) The average CERES score for the DepMap cohort of 20 MM cell lines (x-axis) plotted by the average CSS for the 17 MM cell lines in this study. Genes from Fig. S1A, as well as NRAS, KRAS and SLC3A2, highlighted in red. (C) Identification of KRAS and NRAS dependent MM lines by CSS (y-axis).

**Supplemental Figure 2. Proteomic determination of mutant KRAS and NRAS protein interactomes.** (A) Western blot analysis of BioID2 constructs in MM cell lines. (B) Immunofluorescence of BioID2 constructs in RPMI 8226 and SKMM1 cells with anti-BioID2 (red), anti-RAS (green) and DAPI (blue). Scale bar is 10µm. (C) Venn diagram depicting the total size of the BioID2-KRAS$^{G12V}$ and BioID2-NRAS$^{G12V}$ interactomes in MM cell lines, as described in main text. Canonical RAS effectors are listed. (D) Gene Ontology pathway enrichment of the shared KRAS/NRAS (≥2.0 log2fc) interactome, with inverse log10 P-values with Bonferroni correction plotted on the x-axis.

**Supplemental Figure 3. SLC3A2 and RAS associate and regulate mTORC1.** (A) Combined PLA and Immunofluorescence imaging of SLC3A2 (green) and RAS (red) in RPMI 8226 and SKMM1 cells with PLA (pink) and DAPI (blue) staining. Scale bar is 10µm. Representative images from 2 independent experiments. (B) FACS analysis of surface SLC3A2 (CD98) on MM lines expressing indicated control or RAS-targeted shRNAs. Representative data from 3 independent experiments.

**Supplemental Figure 4. RAS and MTOR associate in MM cells.** (A) Western blot analysis of mTORC1 signaling outputs via p-4EBP1 (S65) and p-p70S6K (T389) following knockdown of KRAS, NRAS or KRAS/NRAS in LP1 and KMS12PE cells expressing wild type RAS isoforms. Representative blots from 4 independent experiments. (B) The average MTOR-RAS PLA puncta/cell for 16 MM cell lines. Each dot represents an individual MM line. P-value from one-tailed Mann-Whitney test; representative data from 2 independent experiments. (C) Correlation between MTOR-RAS PLA (x-axis) and SLC3A2-RAS PLA (y-axis). Spearman r value and P value calculated by correlation analysis in Graphpad. (D) Correlation between MTOR-RAS PLA (x-axis) and SLC3A2 CSS (y-axis). Spearman r value and P value calculated by correlation analysis in Graphpad.

**Supplemental Figure 5. RPTOR regulates SLC3A2-RAS association.** (A) SLC3A2-RAS PLA for RPMI 8226 and SKMM1 cells expressing control shRNAs or shRNAs targeting RPTOR or RICTOR. *** denotes p-value <0.0001, **=0.0024 by one-way ANOVA. Scale bar is 10μm. Data from 3 independent experiments. (B) Imaging of RPTOR-RAS or RICTOR-RAS PLA in RPMI 8226 and SKMM1 cells with PLA (red), WGA (green) and DAPI (blue) staining. Scale bar is 10μm. Representative images from 3 independent experiments.

**Supplemental Figure 6. A large-scale combinatorial drug screen reveals synergistic interaction and druggable vulnerabilities of RAS-dependent MM lines.** Drug Set Enrichment Analysis (DSEA) of the Everolimus vs MIPE5.0 screen in SKMM1 and PRMI 8226. The average Excess HSA was used to pre-rank combinatorial outcomes before running DSEA. Enrichment plots for MEK (A) and ERK (B) inhibitors are shown, together with the 6x6 blocks for the 3 most-synergistic drugs within each target-class.

**Supplemental Figure 7. Combinations of mTORC1 and MEK inhibitors induce apoptosis in MM cells.** (A) Analysis of cell cycle and viability in XG2 (left) and L363 (right) MM cell lines treated with DMSO, everolimus, trametinib or both drugs. For Dyecycle cell cycle analysis, bars are labeled G1, S or G2 with indicated percentages. For 7AAD/Annexin V and cleaved caspase-3 staining percent of cells in indicated gates are listed in plots. Representative experiment from at least 2 independent experiments. (B) Western blot analysis of MM lines treated with indicated drugs. Representative blots from 3 independent experiments.

**Supplemental Figure 1**

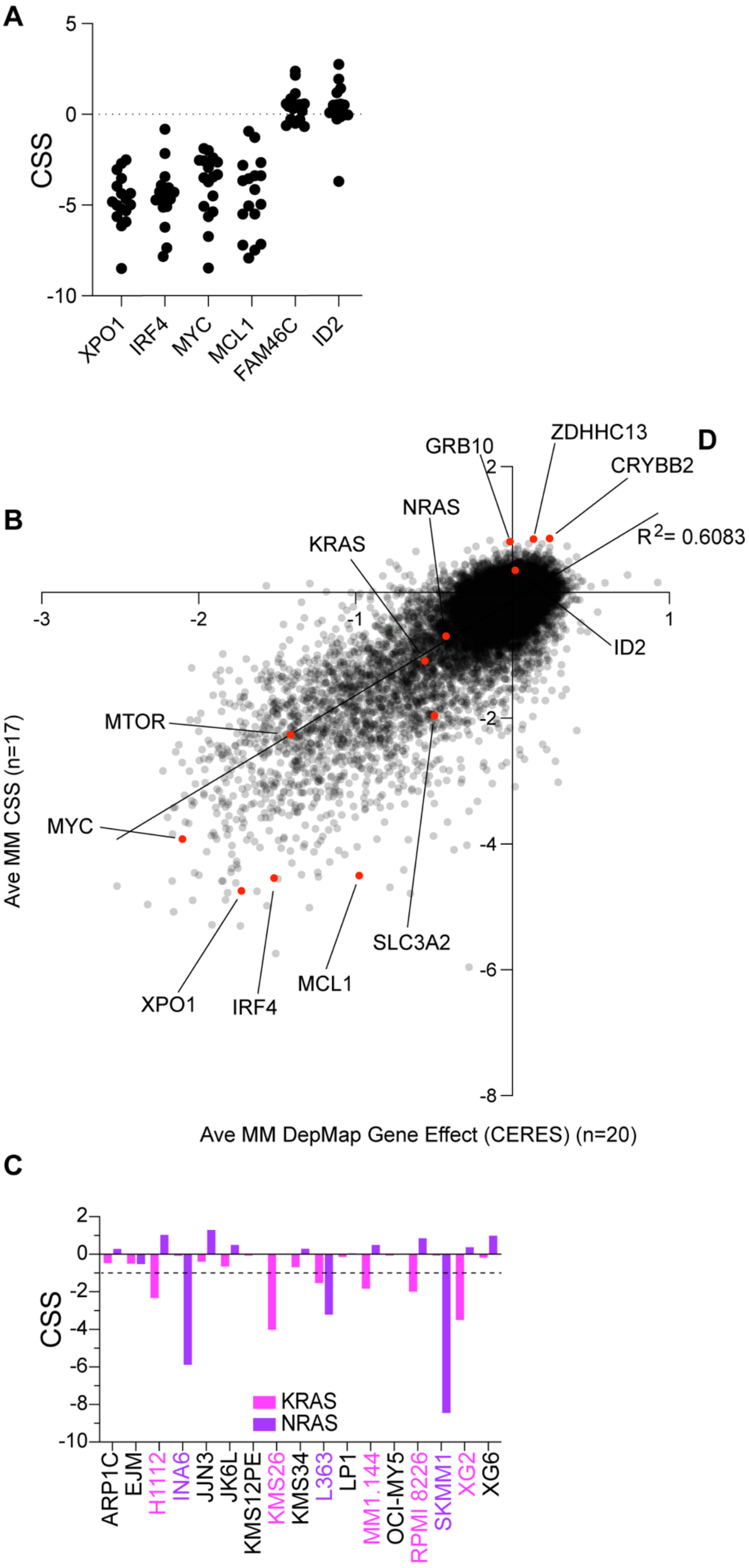

**Supplemental Figure 2**

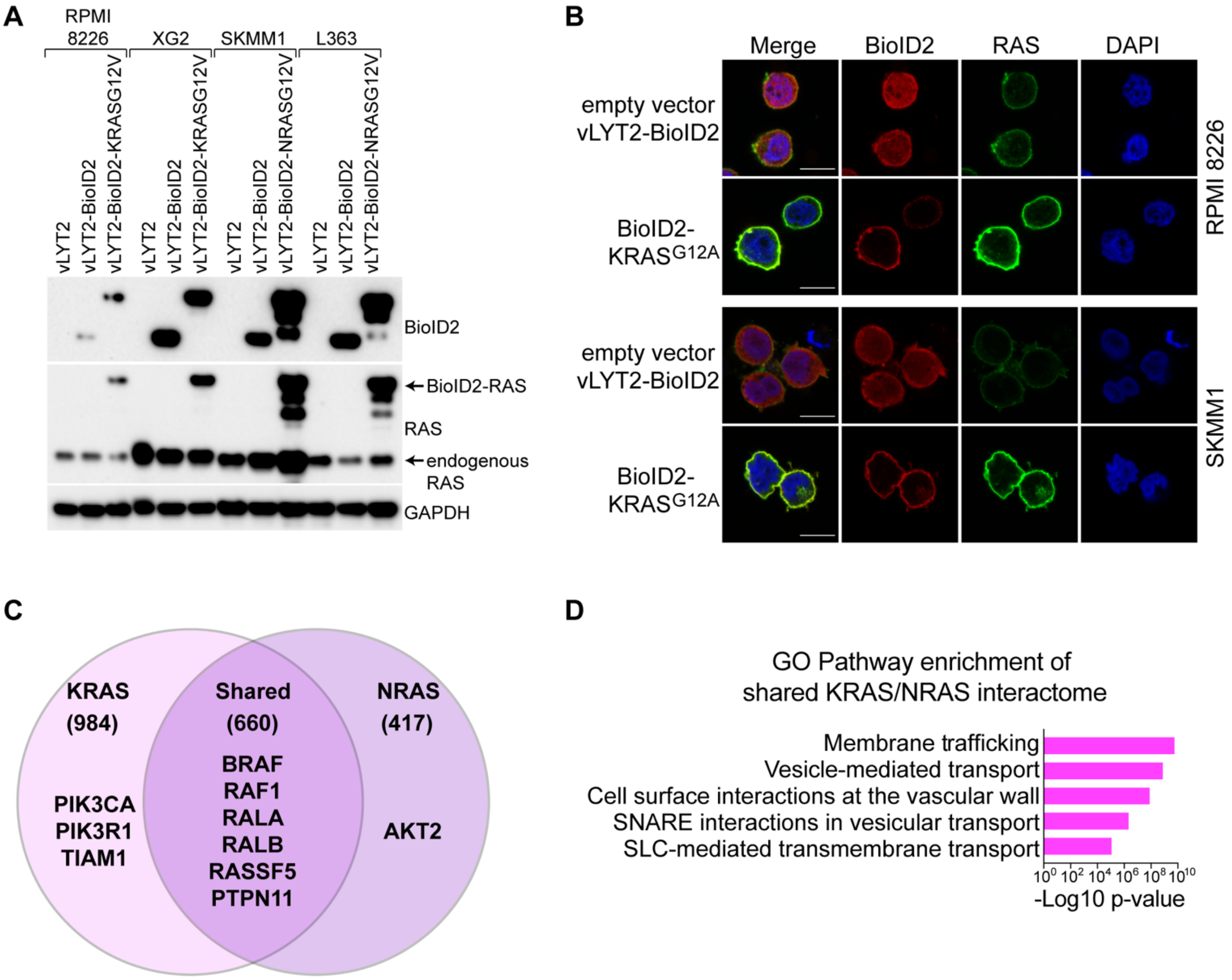

### Supplemental Figure 3

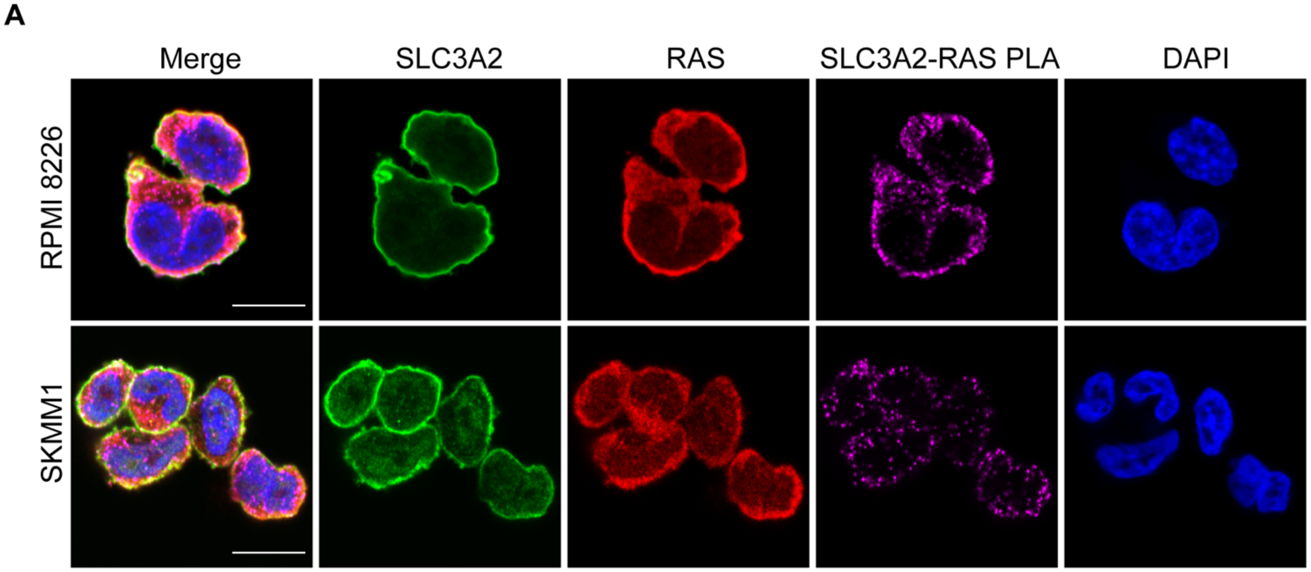


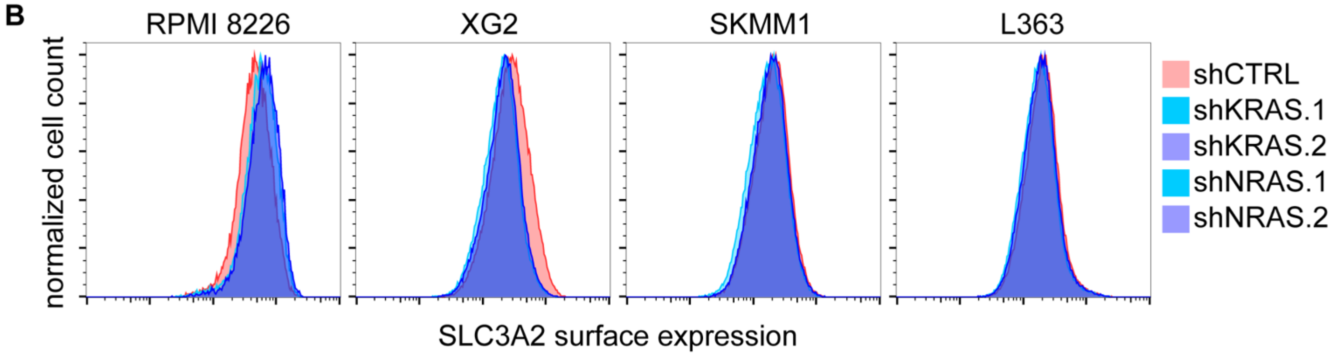

### Supplemental Figure 4

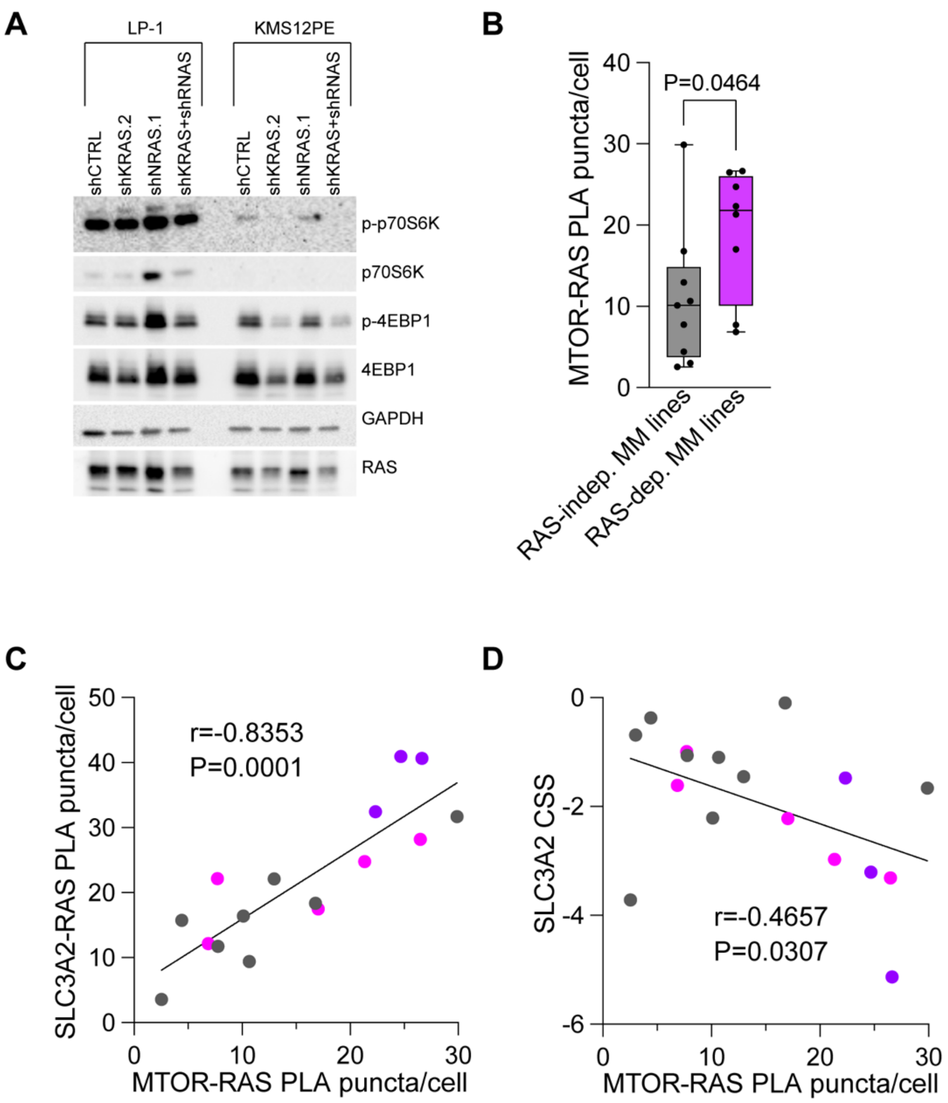

**Supplemental Figure 5**

A

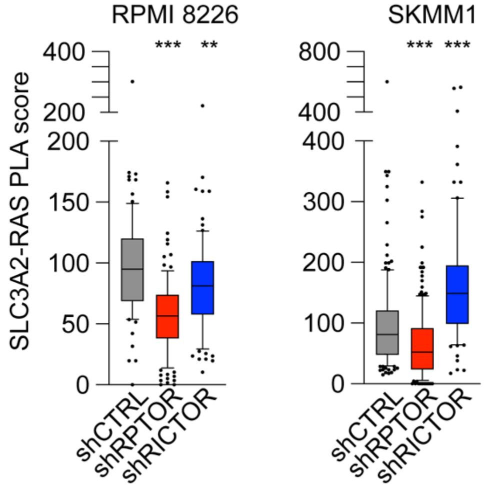


B

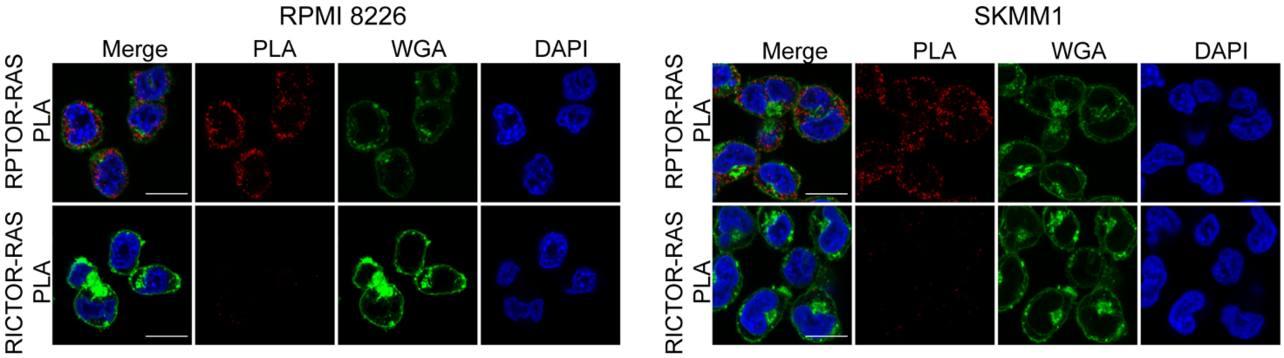

### Supplemental Figure 6

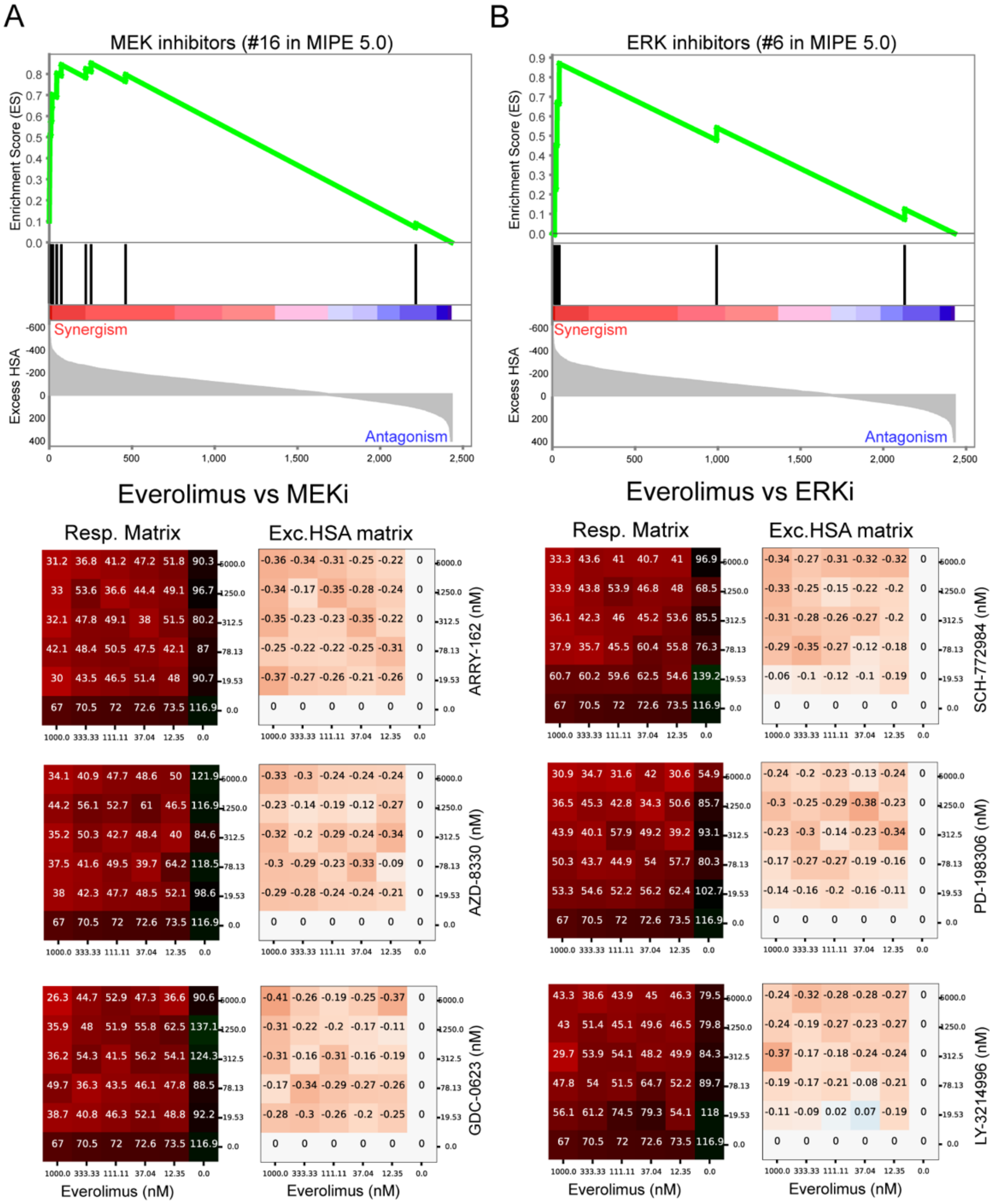

### Supplemental Figure 7

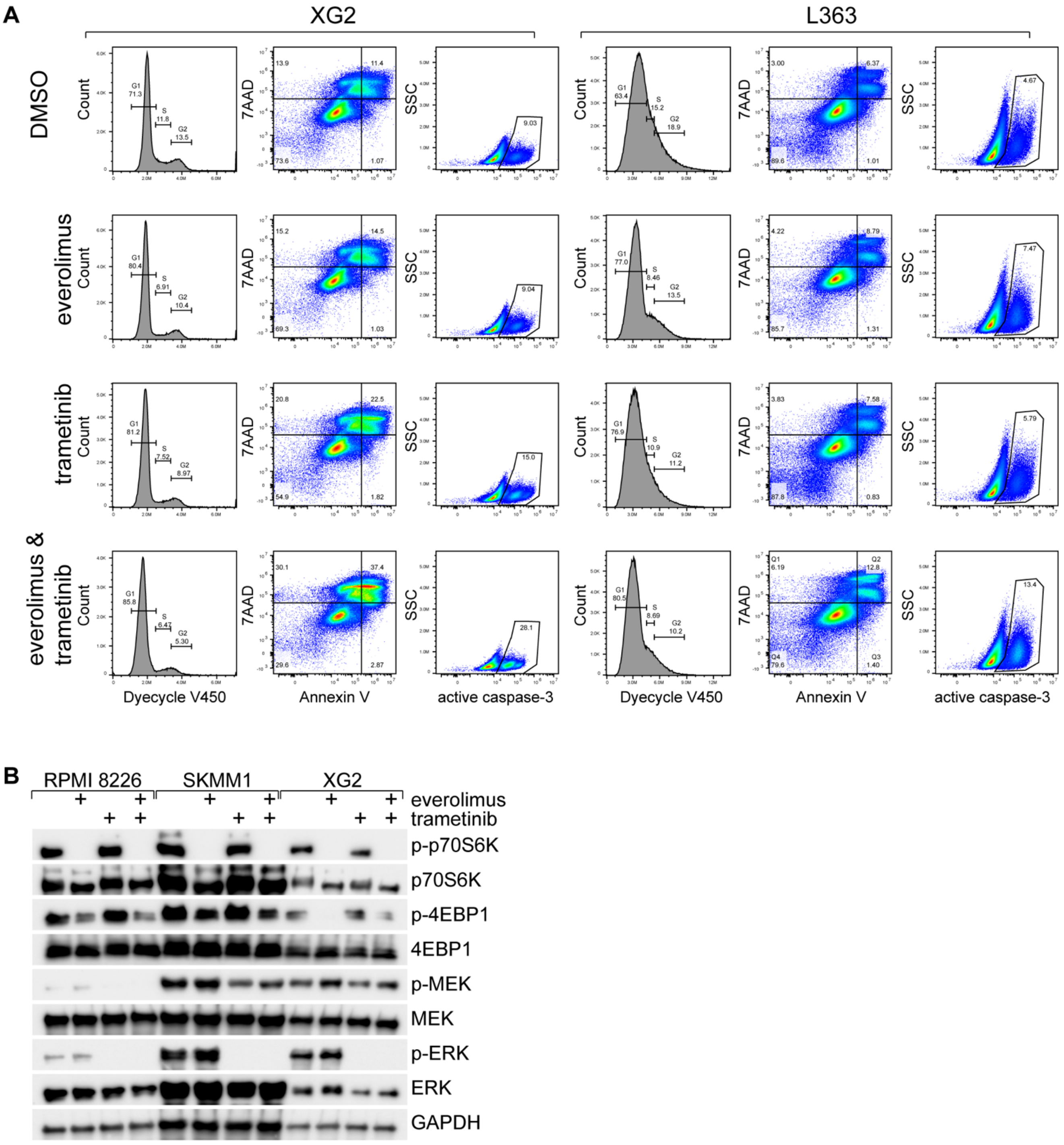
